# The Arp2/3 complex maintains genome integrity and survival of epidermal Langerhans cells

**DOI:** 10.1101/2024.12.27.630538

**Authors:** Maria-Graciela Delgado, Ann-Kathrin Burkhart, Charlotte Canet-Jourdan, Javiera Villar, Roberto Amadio, Vanessa S. Racz, Doriane Sanséau, Nilushi De Silva, Livia Lacerda Mariano, Anna Shmakova, Anna Chipont, Mabel San Roman, Giulia Bracchetti, Vincent Calmettes, Zahraa Alraies, Mathieu Maurin, Aurelien Dauphin, Coralie Guerin, Florent Ginhoux, Nicolas Manel, Daniele Fachinetti, Federica Benvenuti, Ana-Maria Lennon-Duménil, Sandra Iden

**Author notes:** shared first authors (Maria-Graciela Delgado, Ann-Kathrin Burkhart). shared senior authors. **CONTACT DETAILS:** SI: Cell and Developmental Biology; Center for Center for Gender-specific Biology and Medicine (CGBM); Center for Human and Molecular Biology (ZHMB), Faculty of Medicine, Kirrberger Strasse 100, Building 61.4, D-66421 Homburg/Saar, Germany Saarland University, Germany, AML: INSERM U932, Immunity and Cancer, Institut Curie, PSL University, Paris, France.

## Abstract

Myeloid cells use intracellular actin networks for key cellular processes, including cell migration and chemotaxis, phagocytosis or macropinocytosis, as well as immune synapse formation. However, whether actin networks play any role in the development and/or survival of myeloid cells in tissues remains open. Here, we found that the Arp2/3 complex, which is responsible for the nucleation of branched actin networks, is needed for *in vivo* maintenance of epidermal Langerhans cells (LCs) throughout life. Mice harboring a genetic deletion of the Arpc4 subunit of the complex in myeloid cells form LC networks at birth, but these cells decline in numbers following a process reminiscent of premature cellular aging. By combining *in vivo* analyses of LCs with *in vitro* experiments on bone-marrow-derived dendritic cells, we found that Arpc4-deficient cells manage to progress through the cell cycle but accumulate DNA damage associated with aberrant nuclear shapes, lamina reduction and events of nuclear envelope rupture. These results provide the first evidence for a physiological role of Arp2/3 in maintenance of genome integrity and survival of immune cells in tissues.

## INTRODUCTION

The success of immune responses relies on the capacity of myeloid cells such as macrophages and dendritic cells (DCs) to develop in their niches, colonize peripheral tissues where they often complete their differentiation, and eventually migrate from these tissues to lymphoid organs to alert lymphocytes upon invasion by microorganisms. It is well accepted that the migration of myeloid cells is controlled by the actin cytoskeleton (Kamnev et al., 2021), however, the role of this cellular machinery in myeloid cell development and maintenance remains largely unknown.

Distinct subcellular actin structures, such as adhesions and protrusions, are used during development, for instance by embryonic and mesenchymal stem cells, to sense the biochemical and/or mechanical cues present within their environment that guide their differentiation and define their survival (Putra et al., 2023; Sen et al., 2017). A major actor in the formation of such actin structures is the Arp2/3 complex, which is responsible for the nucleation of branched actin networks in all eukaryotic cells. Arp2/3 is composed of seven subunits and is activated by Nucleation Promoting Factors (NPFs) such as WAVE, WASP and WASH, which are regulated by small GTPases of the Rho family, namely Rho, Rac and Cdc42 (Gautreau et al., 2022; James et al., 2024). The complex is further inhibited by Arpin (Dang et al., 2013). Intriguingly, recent studies focusing on Arpin have highlighted several new roles for Arp2/3 in regulation of the cell cycle (Molinie et al., 2019) and maintenance of DNA integrity (Simanov et al., 2024). For instance, nucleation of branched actin by Arp2/3 has been shown to help assemble DNA double-strand breaks for repair (Schrank et al., 2018; Caridi et al., 2018; Zagelbaum et al., 2023). Moreover, Arp2/3 was found to promote the maintenance and remodeling of replication forks (Lamm et al., 2020; Palumbieri et al., 2023; Nieminuszczy et al., 2023). However, whether these observations so far made in cultured cells have any physiological relevance *in vivo* remains an open question.

Langerhans cells (LCs) are epidermis-resident myeloid cells that belong to the macrophage lineage but share migratory properties with DCs (Ginhoux et al., 2006; Hoeffel et al., 2012; Kaplan 2017; Doebel et al., 2017). LCs form a tissue-wide network within the epidermis and act as barrier of defense against external threats (Clausen & Stoitzner, 2015; Deckers et al., 2018). Previous studies have shown that deletion of Rac1 or Cdc42 perturbs the maintenance of epidermal LC networks (Luckashenak et al., 2013; Park et al. 2021). However, the mechanisms resulting in these defects remain elusive, prompting us to investigate the role of Arp2/3 in LCs. We found that while Arp2/3 is not needed for the early postnatal establishment of LC networks, LCs lacking the ArpC4 subunit of the complex progressively decline with age. This is associated with nuclei exhibiting aberrant shapes and displaying hallmarks of premature cellular aging: decreased levels of Lamin A/C and B1, loss of nuclear envelope integrity and accumulation of DNA damage. Using bone-marrow-derived DCs as a cellular model, we found that this phenotype results at least in part from defects in DNA repair in proliferative differentiating cells. Accordingly, assessing different myeloid populations *in vivo* revealed that proliferating cells, such as LCs and alveolar macrophages, but not quiescent myeloid cells, require Arp2/3 for their maintenance. Collectively, our results provide evidence for a physiological role of Arp2/3 in nuclear shape, genome integrity and aging of myeloid cells in specific peripheral tissues.

## RESULTS

### ArpC4 gene deletion results in reduced Langerhans cell numbers in the mouse epidermis

To examine a potential role of Arp2/3 in LC networks, we crossed mice harboring a loxP-flanked allele for the *Arpc4* essential unit of this complex (see methods; van der Kammen et al., 2017) with CD11c-Cre animals (Caton et al., 2007) (*CD11cCre*;*Arpc4*^fl/fl^, hereafter referred to as *Arpc4*^KO^). CD11c is known to be expressed by LCs around postnatal day 2 to 4, after they have entered the epidermis (Chorro et al., 2009), and by precursors of DCs that reside in the dermis (Clausen & Stoitzner, 2015). *Arpc4* gene deletion was verified at the protein level in bone-marrow-derived CD11c^+^ DC cultures (Rivera et al., 2022). To assess the consequence of Arpc4 loss for LCs, we first analyzed whole-mount epidermis from ear skin of adult 8-10 week-old control and *Arpc4* mutant mice. Strikingly, we noted a significant decrease of LC numbers in the ears of *Arpc4*^KO^ mice (Fig. 1A,B). Voronoi diagrams further served to define the distance between neighboring LCs and the area each LC occupied (Fig. 1A,C; Fig. S1A), revealing a globally disrupted LC network upon *Arpc4* loss with mutant LCs being more isolated than their wild-type counterparts. The decreased LC count of *Arpc4*^KO^ mice was further confirmed by flow cytometry analysis of the LC population in ear skin (Fig. 1D; Fig. S1B). In addition, Arpc4-deficient LCs were characterized by an altered LC morphology with reduced dendritic complexity (Fig. 1E-G). The decline of LCs was also evident when analyzing the recently reported stereotypical LC pattern observed in the mouse tail epidermis (Baess et al., 2022) (Fig. 1A-C,E,G). Collectively, these results suggest an essential role for Arp2/3 in the homeostasis and morphology of epidermal LCs.

**Figure 1:**
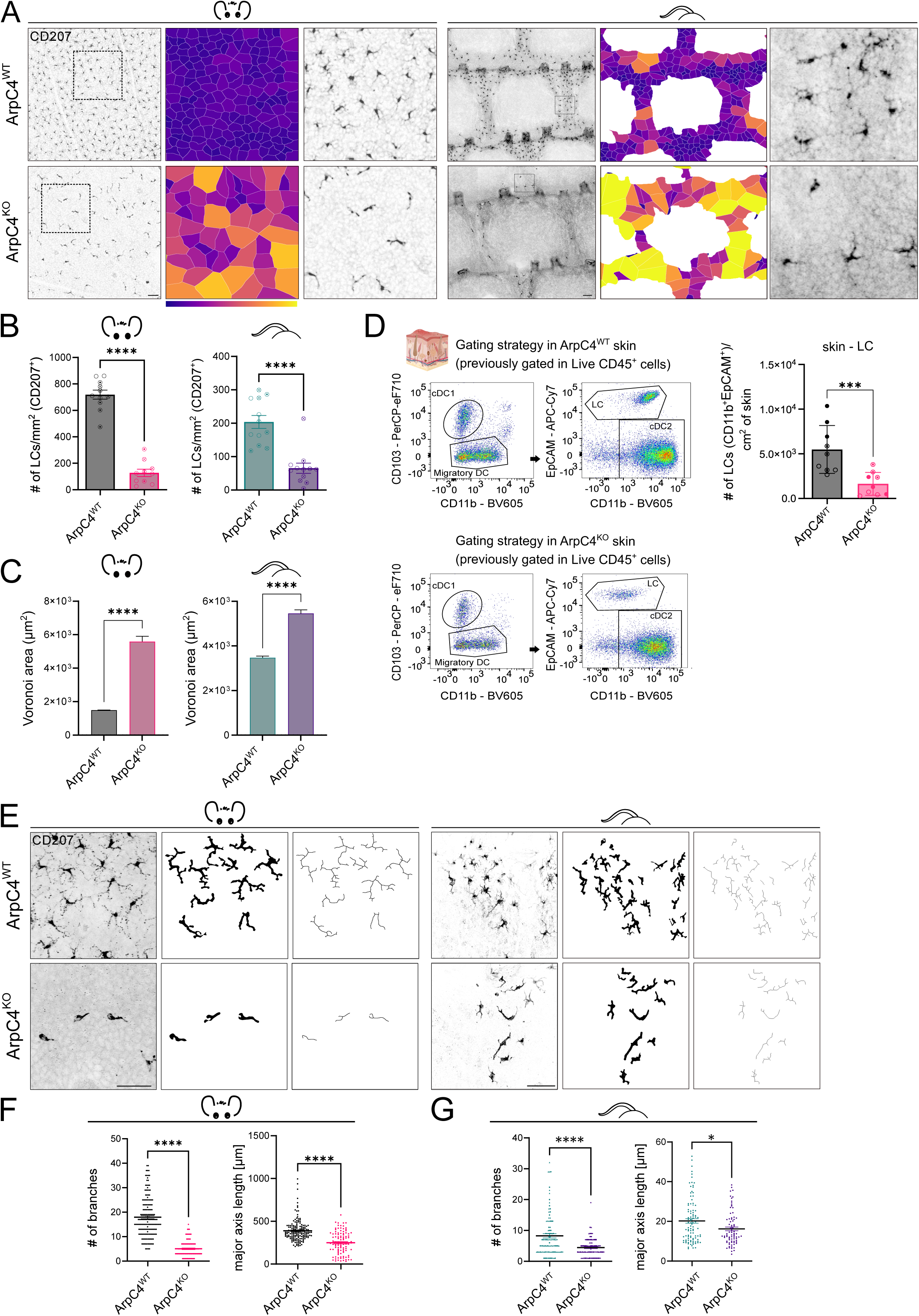
The number of LCs is strongly decreased in the ear and tail epidermis of conditional *Arpc4*^KO^ mice. (A) Micrographs of ear (left panel) and tail (right panel) epidermal sheets from adult (8-12 weeks old) control (ArpC4^WT^) and ArpC4^KO^ mice immunostained for LC marker (CD207/langerin). Middle: Voronoi diagram of LCs generated from micrographs, color scale bar 0 mm^2^*=*purple – 14 mm^2^= yellow. Right: magnification of marked areas. Scale bars = 100 µm. (B) Quantification of (A): LC density (LC number/mm²) in ear (left) and tail (right) epidermis (n = 12 mice; ****p = <0.0001; mean ± SD; unpaired two-tailed Student’s t-test). Matching symbols denote animals used in the same experiment. (C) Quantification of area occupied by LCs from Voronoi diagrams in ear (left) and tail (right) epidermis (n= 6 mice, 3 ctrl /3KO, n ear = 2016 ctrl, 599 KO cells; n tail = 1848 ctrl, 1072 KO cells, ****p = <0.0001; mean ± SD; unpaired two-tailed Student’s t-test with Welch’s correction. (D) Gating strategy in skin of ctrl (upper panel) and KO (lower panel), gates on cDC1 and DC migratory (left) were used to identify LC and cDC2. Right panel: LC numbers in the skin (CD11c^+^, EpCAM^+^) (N=9, 3 mice per experiment, 3 experiments, mice aged 8–14 weeks. ***p = 0.0008; mean ± SD; two-tailed Mann-Whitney U-test). Matching symbols denote animals used in the same experiment. Skin scheme from Biorender (www.biorender.com). (E) Skeletonized images of micrographs of ear (left panel) and tail (right panel) epidermal sheets of 8–12-week-old mice immunostained for CD207/langerin. (F) Quantification of E (left panel): dendricity of LCs (branches/LC) (left) and major axis length (right) in ear epidermis. (N = 3 mice; n = 686 ctrl, 285 KO cells; ****p<0.0001, ***p = 0.0007; mean ± SD; two-sided Mann-Whitney U-test) (G) Quantification of E (right panel): dendricity of LCs (branches / LC) (left) and major axis length (right) in tail epidermis. (N = 3 ctrl, 4 KO mice; n = 99 ctrl, 80 KO cells; ****p<0.0001, *p = 0.0271; mean ± SD; two-sided Mann-Whitney U-test). Scale bars = 50µm. WT=wildtype, KO=knockout

### Arpc4 deficiency alters LC networks in adolescent mice

It has been shown that LCs develop from fetal liver embryonic precursors (CX3CR1^+^ CD45^+^ Langerin^−^) that colonize the epidermis around birth, a time at which they start expressing MHC class II, CD11c and, slightly later, Langerin (Tripp et al., 2004; Chorro et al., 2009). LCs from *Arpc4*^KO^ mice should therefore delete the *Arpc4* gene at this precursor stage. Once spread within the epidermis, these cells undergo a massive burst of proliferation that leads to formation of an intricate LC network at P7, whereas only ∼1-8% of LCs are proliferating to maintain LC homeostasis in adult mice (Merad et al., 2008; Chorro et al., 2009; Ghigo et al., 2013; Zaru et al., 2015). To assess whether the decline of LCs observed in *Arpc4*^KO^ mice was due to defects in epidermal homing and early postnatal expansion, we analyzed LC networks during the first week of mouse post-embryonic development. While the proliferative LC burst was evident when comparing P1 and P7 skin, we found that the numbers of LCs did not differ between control and *Arpc4*^KO^ ear and tail epidermis (Fig. 2A,B). In line with this result, we did not detect significantly altered proliferation or apoptosis of LCs in P7 *Arpc4*^KO^ epidermal sheets, as assessed by immunohistochemistry for Ki67 (Fig. 2C,D) and cleaved Caspase 3 (Fig. S2A,B), respectively. Importantly, Cre-mediated recombination of the *Arpc4* locus was verified by PCR of P7 epidermal tissue samples (Fig. S2C), confirming successful CD11cCre-mediated gene deletion. Together, these data show that the strongly reduced numbers of LCs in the epidermis of adult *Arpc4*^KO^ mice do not result from impaired initial formation of the LC network from precursor cells and instead suggest that Arp2/3 is required for subsequent LC network maintenance.

**Figure 2:**
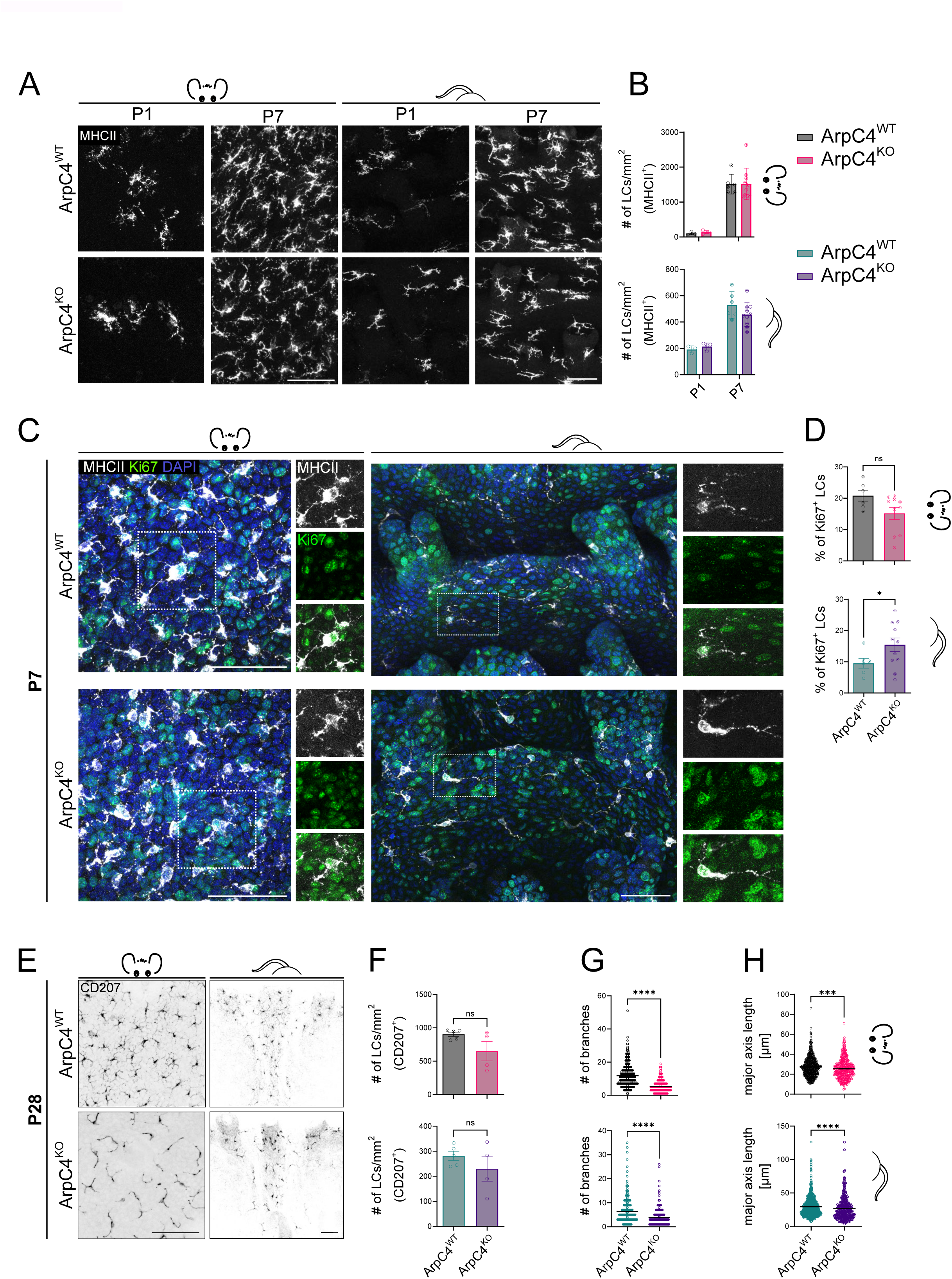
Deletion of Arpc4 does not interfere with the initial establishment of postnatal LC networks. (A) Micrographs of ear (left) and tail (right) epidermal sheets of P1 and P7 LC *ArpC4*^WT^ and *ArpC4*^KO^ mice stained for early LC marker (MHCII). (B) Quantification of (A): LC density (LC number/mm²) in ear (top) and tail (bottom) epidermis. Ear P1: (n = 3 ctrl, 4 KO mice); P7: (n = 6 ctrl, 10 KO mice), Tail P1: (n = 3 ctrl, 4 KO mice); P7: (n = 6 ctrl, 10 KO mice), Two-way ANOVA/Tukey’s multiple t-test; mean ± SD (C) Micrographs of LC (MHCII^+^; grey) and Ki67 immunostaining (green) in P7 ear (left) and tail (right) epidermal sheets. DAPI (blue) serves as nuclear counterstain. (D) Quantification of (C): percentage of Ki67-positive LCs in ear (top) and tail (bottom) epidermis at P7. Ear: (n = 6 ctrl, 10 KO mice; ns p = 0.1806; mean ± SD; two-sided Mann-Whitney U-test). Tail: (n = 6 ctrl, 11 KO mice; *p = 0.0422; mean ± SD; Welch test). (E) Micrographs of ear (left) and tail (right) epidermal sheets of 4-weeks-old LC *ArpC4*^WT^ and *ArpC4*^KO^ mice immunostained of LC marker (CD207/langerin). (F) Quantification of (E): LC density (LC number/mm^2^) in top: ear and bottom: tail epidermis. Ear: (N = 5 ctrl, 4 KO mice; ns p = 0.1905; mean ± SD; two-sided Mann-Whitney U-test). Tail: (N = 5 ctrl, 4 KO mice; ns p = 0.3264; mean ± SD; unpaired, two-tailed t-test). (G) Quantification of (E): dendricity of LCs (branches / LC) in ear (top) and tail (bottom) epidermis. Ear: (N = 5 mice; n = 578 ctrl, KO cells, 485 KO cells; ****p<0.0001; mean ± SD; Two-sided Mann-Whitney U-test); Tail: (N = 5 ctrl, 4 KO mice; n = 624 ctrl, 371 KO cells; ****p<0.0001; mean ± SD; two-sided Mann-Whitney U-test). (H) Quantification of (E): major axis length of LCs in ear (top) and tail (bottom) epidermis Ear: (N = 5 mice; n = 578 ctrl, 485 KO cells; ***p = 0.0002; mean ± SD; Two-sided Mann-Whitney U-test; Tail: (N = 5 ctrl, 4 KO mice; n = 624 ctrl, 371 KO cells; ****p<0.0001; mean ± SD; two-sided Mann-Whitney U-test). Scale bars = 50µm

To unravel the underlying mechanisms, we first attempted to better define the postnatal kinetics followed by these cells. Imaging epidermal sheets from 4 weeks-old mice showed that the number of LCs started to decrease in some *Arpc4*^KO^ samples even though this difference did not yet reach statistical significance (Fig. 2E,F). Similarly, a trend for increased apoptosis was observed in ear and tail skin (Fig. S2D,E). We therefore chose the 4-week time-point for further investigation of possible molecular and cellular effects of ArpC4 deletion, as at this stage the loss of LCs first became apparent. Strikingly, when analyzing the morphology of these cells, we found that *Arpc4*^KO^ LCs were less elongated and exhibited less dendrites as compared to their wild-type counterparts (Fig. 2G,H; Fig. S2F), just as observed in adult mice (Fig. 1E-G), indicating that *Arpc4* deficiency already impacted the morphology of LCs at this time-point. These results suggest that LCs start dying in adolescent *Arpc4*^KO^ mice.

### Arpc4 loss elicits nuclear aberrations in bone-marrow-derived DCs and in epidermal LCs

To gain further insights into the molecular and cellular mechanisms underlying the premature decline of LCs in *Arpc4*^KO^ LCs, we searched for an *ex vivo* culture model that allows experimental manipulation. Unfortunately, at present there is no suited model for the study of murine LCs that would sufficiently recapitulate LC functions. We thus turned to bone-marrow derived DCs, which share important features with LCs: they (1) display a dendritic morphology, (2) ingest extracellular material, (3) rely on cell-cell interactions to differentiate *ex vivo* and grow in clusters (Inaba et al., 1992; Inaba et al. 2009), and (4) acquire a migratory phenotype in response to inflammation (Matzner et al., 2008). We found that control and *Arpc4*^KO^ DCs differentiated from bone-marrow precursors in the presence of GM-CSF were phenotypically similar in terms of MHC class II and CD86 expression (Fig. S3A-C), indicating no major developmental and maturation defects in the absence of Arp2/3.

Unlike stable cell lines, these primary DC cultures display some degree of heterogeneity e.g. in marker gene expression and cell morphology. To handle the variable cell shapes seen in DCs, we chose to image cells after introducing them in microchannels, which helps normalizing their shape and quantifying organellar morphologies (Vargas et al., 2016). Noticeably, we found that *Arpc4*^KO^ DCs displayed abnormal nuclear shapes, with deformed, two- or multilobular nuclei of different sizes, sometimes connected by DNA bridges (Fig. 3A; Fig.3B, red arrows). These aberrantly shaped nuclei could also be observed by electron microscopy (Fig. S3D,E). In addition, staining for the nucleoskeleton components Lamin (Lmn) A/C and Lmn B1 showed a significant decrease in *Arpc4*^KO^ DCs, no difference being observed when staining their nuclear envelope with Lap2b antibodies (Fig. 3C,D). Importantly, such reduction of Lmn A/C and B1 immunoreactivity could be recapitulated in LCs from epidermal sheets *in vivo* (Fig. 3E,F). Similarly, super-resolution analysis confirmed a significant increase of aberrantly shaped nuclei in LCs from the epidermis of *Arpc4*^KO^ mice (Fig. 3G,H). Immunofluorescence analysis for the Arp2/3 complex member Arpc5 further revealed the presence of a fraction of Arpc5 within the nucleus of DCs (Fig. S3F,G). These data show that both *Arpc4*^KO^ LCs and DCs exhibit major alterations in the shape and skeleton of the nucleus and are suggestive of the presence of the Arp2/3 complex within this organelle.

**Figure 3:**
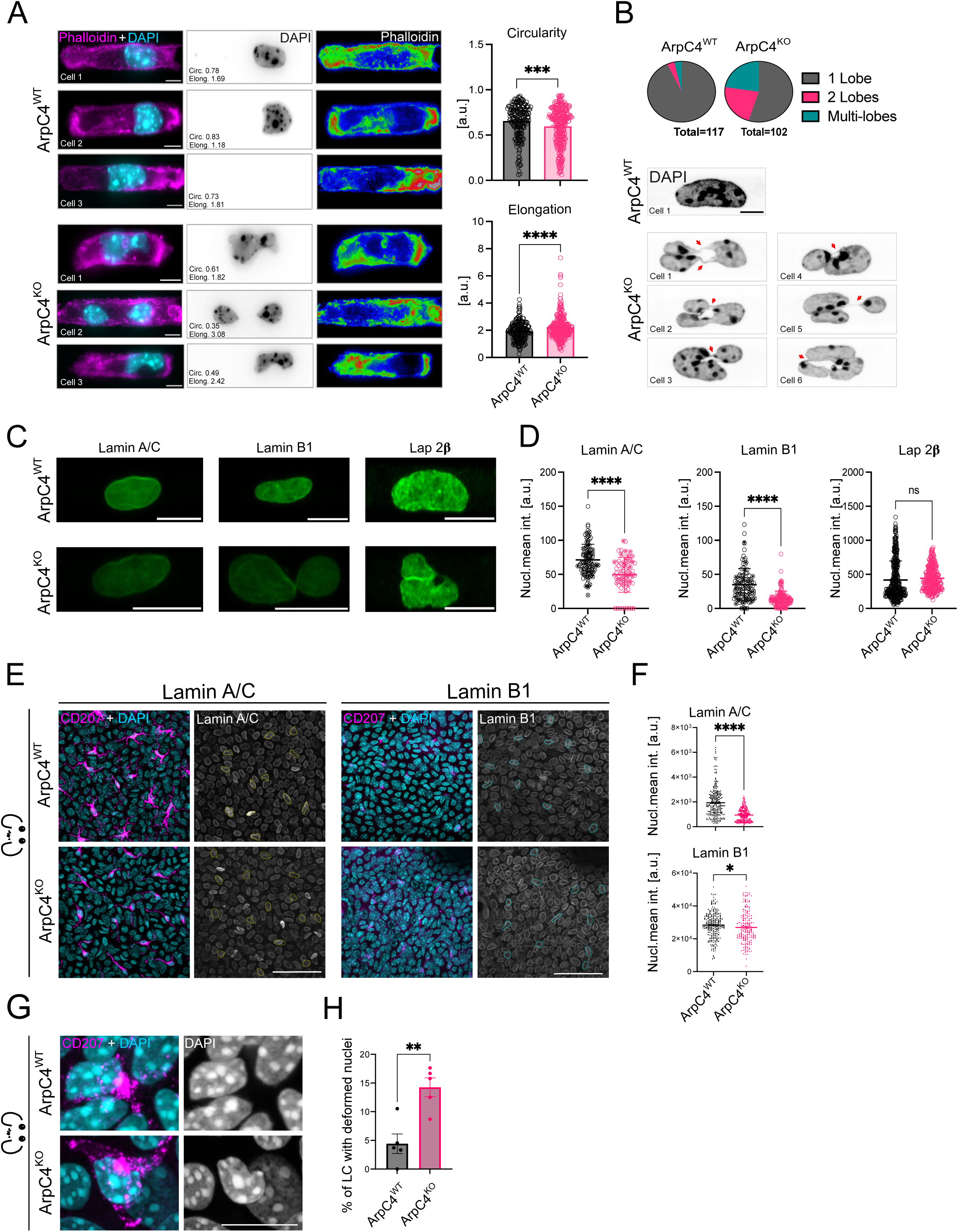
Aberrant nuclear shape in *Arpc4*^KO^ DCs and LCs. (A) Left: Exemplary confocal micrographs of BMDCs inside 8 μm × 5 μm fibronectin-coated µ-channels stained for Phalloidin (magenta) and DAPI (cyan); middle: DAPI (grey), right: Phalloidin density maps, showing actin cytoplasmic distribution. Bar diagrams (right panel) represent circularity and elongation values indicating nuclear shape factors. Scale bars = 6 μm. (circularity: n ctrl= 224, n KO= 302, ***p=0.0009; mean ± SD; unpaired, two-tailed t-test); elongation, n ctrl = 221, n KO= 287, ****p<0.0001; mean ± SD; unpaired, two-sided t-test). (B) Quantification of nuclear lobes in *ArpC4*^WT^ and *ArpC4*^KO^ DCs. Top panel: Pie charts for ctrl (n=117) and KO (n=102) based on manual analysis of the number of lobes/nucleus. Bottom panel: Exemplary OMX-Super resolution images of BMDCs inside 8 μm × 5 μm fibronectin-coated µ-channels showing DAPI signal (grey). Red arrows indicate DNA bridges. Scale bars = 6 μm. (C) Representative Z-stack projection of confocal images of BMDC nuclei inside 8 μm × 5 μm fibronectin-coated µ-channels immunostained for lamin A/C (right), lamin B1 (center), and Lap-2β. Scale bars = 10 μm. (D) Quantification of nuclear mean fluorescence intensity of lamin A/C (n ctrl= 97, n KO= 68, ****p<0.0001; mean ± SD; unpaired, two-sided t-test), lamin B1 (n ctrl= 101, n KO= 115, ****p<0.0001; mean ± SD; unpaired, two-sided t-test), Lap-2β (n ctrl= 396, n KO= 320, ****p<0.0001; mean ± SD; unpaired, two-sided t-test). (E) Micrographs of immunostainings for LCs (CD207^+^; magenta) and lamin B1 (left) or lamin A/C (right) in ear epidermal sheets of 4-weeks-old *ArpC4*^WT^ and *ArpC4*^KO^ mice. Margins of LC nuclei were outlined manually (Lamin B1: blue and Lamin A/C: yellow). DAPI is shown in blue. Scale bar = 50 µm. (F) Quantification of (E). Nuclear signal intensity of lamin B1 (top) and lamin A/C (bottom) immunostaining in ear epidermis LCs. Ear: Lamin B1: (N = 4 ctrl, 5 KO mice; n = 219 ctrl, 179 KO cells; *p = 0.021; mean ± SD; two-sided Mann-Whitney U-test). Lamin A/C: N = 5 mice; n = 241 ctrl, n = 251 KO cells; ****p = <0.0001; mean ± SD; two-sided Mann-Whitney U-test). (G) Micrographs of nuclei (DAPI, blue) and LCs (CD207^+^, magenta) in ear epidermal sheets of 4-weeks-old mice. Scale bar = 10µm. (H) Quantification of (G): percentage of LCs with deformed nuclei. N = 5 ctrl mice, 5 KO mice; n = 65 ctrl, 44 KO cells; **p = 0.0032, mean ± SD, unpaired two-tailed Student’s t-test.

### Arpc4^KO^ DCs exhibit aberrantly shaped, fragile nuclei and increased DNA damage

Decreased Lamin levels have recently been shown to lead to nuclear envelope rupture and premature aging of lung macrophages (De Silva et al., 2023), prompting us to analyze nuclear envelope integrity in *Arpc4*^KO^ DCs. Staining these cells with an antibody against Lap-2β revealed that their aberrantly shaped nuclei occasionally showed discontinuity of the nuclear envelope when imaged by super-resolution microscopy, suggesting that it had lost its integrity (Fig. 4A), accompanied by a decrease in nuclear circularity and an increase in nuclear elongation (Fig. 4B). Accordingly, when expressing in DCs an NLS-GFP construct, which leaks into the cytosol upon nuclear envelope breakage, we found that the frequency of nuclear rupture events was increased in *Arpc4*^KO^ DCs as compared to their wild-type counterparts (Fig. 4B,C). Noticeably, while control DCs displayed nuclear envelope ruptures when introduced in confining microchannels (4µm), *Arpc4*^KO^ DCs showed impaired nuclear integrity even in the absence of such physical constraint (in 8µm channels, Fig. 4C,D). Congruent with above findings, *Arpc4*^KO^ DCs also showed increased cytosolic double-stranded DNA (dsDNA) (Fig. 4E,F), further demonstrating that loss of ArpC4 compromises nuclear integrity. Together, these results indicate that Arp2/3 deficiency not only leads to aberrant nuclear shapes and Lamin decrease, but also further fragilizes the nuclear envelope of DCs.

**Figure 4:**
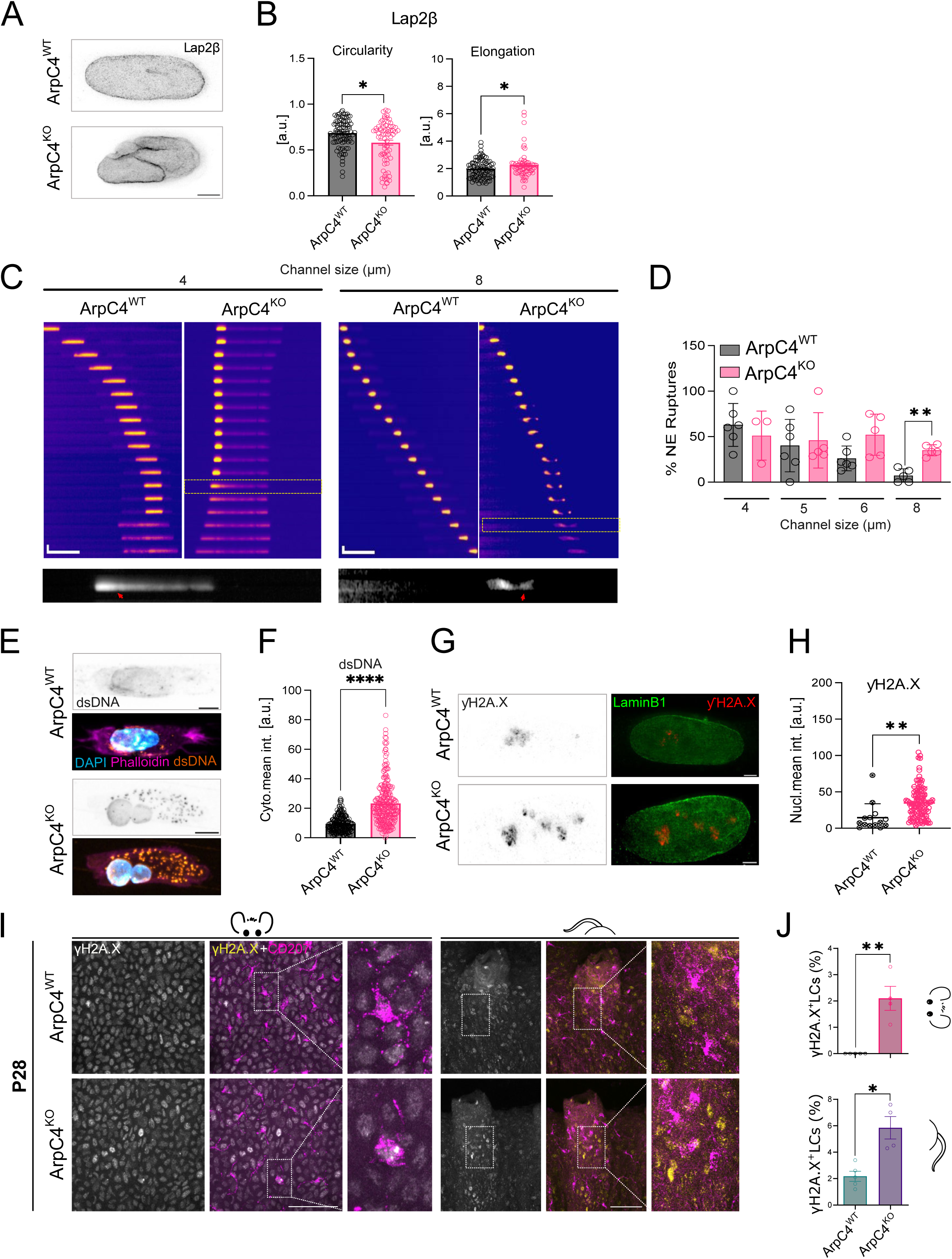
Arp2/3 inhibition leads to aberrantly shaped nuclei characterized by nuclear envelope rupture, DNA damage and signs of premature aging of LCs. (A) Exemplary super-resolution STED images of BMDCs inside 8 μm × 5 μm fibronectin-coated µchannels immunostained for Lap-2β; ctrl (upper panel), KO (bottom |panel). Scale bar = 2µm. (B) Quantification of (A). Bar diagrams represent circularity (left panel) and elongation (right panel) values indicating nuclear shape factors (for circularity: n ctrl = 183, n KO = 186, *p = 0.0210; mean ± SD; unpaired, two-tailed t-test); for elongation: n ctrl = 182, n KO= 181, *p = 0.0253; mean ± SD; unpaired, two-sided t-test). (C) Sequential images of an NLS-GFP positive BMDC (day 7) passing through a 4 μm (left kymographs) and 8 μm (right kymographs) width microchannel from ctrl and KO BMDCs (scale bar width = 50 μm, height= 10 μm), images were acquired every 1 min. Bottom inset shows a magnification of NE rupture, showing NLS-GFP in the cytoplasm (red arrows). (D) Quantification of NE ruptures in BMDCs passing through different channel sizes (4, 5, 6 and 8 μm); ruptured nucleus as seen by GFP signal in NLS-GFP-positive BMDC was quantified and divided by total of nuclei recorded in each condition. (n= 3 cultures/ 4 mice; 2 Ctrl and 2 KO, total nuclei at 4 μm; 28 in Ctrl, 17 in KO. At 5 μm; 18 in Ctrl, 40 in KO. At 6 μm; 28 in Ctrl, 28 in KO. At 8 μm; 6 in Ctrl, 27 in KO. **p = 0.0027; mean ± SD; unpaired, two-sided t-test). (E) Exemplary confocal micrographs of BMDCs inside 8 μm × 5 μm fibronectin-coated µ-channels stained for DAPI (cyan), phalloidin (magenta) and dsDNA (orange), for ArpC4^WT^ (upper panels) and ArpC4^KO^ (lower panels). Scale bar = 5 μm. (F) Quantification of cytoplasmic mean intensity signal for dsDNA in ArpC4^WT^ and ArpC4^KO^ BMDCs (n ctrl= 227, n KO= 257, ****p<0.0001; mean ± SD; unpaired, two-tailed t-test). (G) Representative STED images of BMDC nuclei inside 8 μm × 5 μm fibronectin-coated µchannels immunostained for γH2Ax (left panel, grey), right: merge of Lamin B1 (green) and γH2Ax (red) for both genotypes. The examples shown are from the same micrographs as displayed in A. Scale bar = 2 μm. (H) Quantification of nuclear mean fluorescence intensity of γH2Ax (N= 3 experiments/ 4 mice; 2 Ctrl and 2 KO/experiment, total BMDC; n ctrl= 14, n KO= 107, **p=0.0018; mean ± SD; unpaired, two-sided t-test). (I) Micrographs of immunostainings for LCs (CD207^+^; magenta) and γH2Ax (yellow) in ear (left) and tail (right) epidermal sheets of 4-weeks-old mice. (J) Quantification of (F): percentage of γH2Ax-positive LCs in ear (top) and tail (bottom) epidermis. (Ear: N= 5 ctrl, 4 KO mice; **p = 0.0079; mean ± SD; two-sided Mann-Whitney U-test). (Tail: N = 5 ctrl, 4 KO mice; *p = 0.0151; mean ± SD; unpaired Student’s t-test, two-tailed). Scale bars = 50µm.

In various cell types, it has been observed that loss of Lamins and nuclear envelope integrity facilitates the entry of cytosolic endonucleases into the nucleus and results in DNA damage (Raab et al., 2016), which can promote premature cellular aging (De Silva et al., 2023). Consistent with these observations, we found that *Arpc4*^KO^ DCs displayed increased DNA damage as shown by enhanced staining with an anti-γH2Ax antibody that marks DNA double-strand breaks (Fig. 4G,H). Remarkably, we observed *in vivo* that already at an age of 4 weeks, mice lacking Arpc4 exhibited an increased number of γH2Ax-positive LCs both in ear and tail epidermis (Fig. 4I,J). At 6 weeks, this phenotype was still observed as a trend (Fig. S4A,B), whereas no enrichment of apoptotic LCs was detected at this age (Fig. S4C,D). Of note, comparison of Cd11c-Cre-positive and -negative mice ruled out that the increased DNA damage was merely a consequence of Cre expression (Fig. S4E,F). Together, these results show that Arp2/3 is needed for the maintenance of genome integrity in both DCs and LCs and further suggest that *Arpc4*^KO^ LCs decline in adolescent mice most likely due to accumulation of cellular features associated with premature cellular aging.

### Transcriptomic analysis of ArpC4-deficient DCs reveals altered expression of interferon response and cell cycle-associated genes

We next asked whether the nuclear defects observed in *Arpc4*^KO^ DCs emerged in fully differentiated cells or rather progressively accumulated during differentiation. Indeed, we had previously analyzed the impact of deleting the *Arpc2* subunit of Arp2/3 in fully differentiated DCs without detecting cells with multi-lobulated nuclei (Vargas et al., 2016). In addition, we found that pharmacological Arp2/3 inhibition with CK666 promoted significant γH2Ax nuclear accumulation when added to DCs during differentiation (from day 7 to day 10), but not when treating fully differentiated day 10-cells (Fig. 5A). Consistent with these observations, day 8 *Arpc4*^KO^ DCs also displayed a significant increase in γH2Ax nuclear staining (Fig. 5B). These results prompted us investigating the potential role of Arp2/3 in DC differentiation. To this end, we used bulk mRNA sequencing (RNAseq) to describe the global changes taking place during the differentiation of control and *Arpc4*^KO^ DCs. Cells were analyzed at days 7, 8, 9 and 10 of culture, day 10 corresponding to the fully differentiated state. We observed that surface expression of the CD11c marker, which drives *Arpc4* deletion in our mouse model, gradually increased during differentiation of both control and *Arpc4*^KO^ DCs, no difference being observed between the two genotypes (Fig. 5C). We performed a differential gene expression analysis using DESeq2 comparing control and *Arpc4*^KO^ DCs at each time point. Principal Component Analysis (PCA) showed that the three samples of control DCs grouped by day displayed limited dispersion as compared to *Arpc4*^KO^ DC samples (Fig. 5D, black versus pink symbols), pointing to a more pronounced transcriptomic heterogeneity in differentiating *Arpc4*^KO^ DCs. Noticeably, assessment of *Arpc4* gene expression highlighted that it transiently increased in control cells (Fig. S5A), reaching a peak at day 8. This observation was consistent with Arp2/3 activity exerting a stronger effect during DC differentiation (days 7 to 9) than on fully developed cells, as hypothesized.

**Figure 5:**
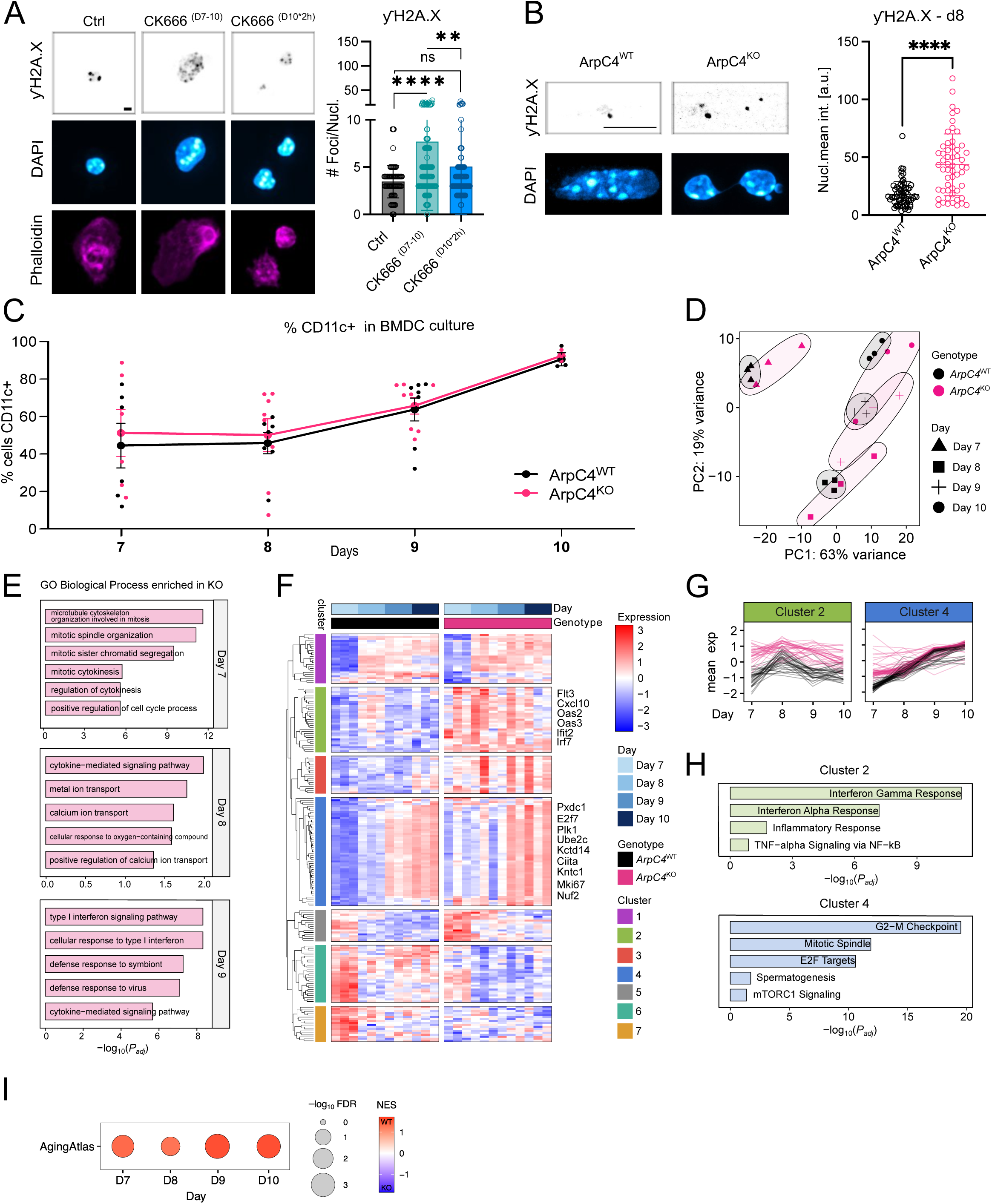
Arpc4 loss-of-function affects DC nuclear integrity, DC transcription and differentiation. (A) Left: Representative confocal images of C57BL6/J BMDC adhered to fibronectin-coated glass coverslip immunostained for γH2Ax, or stained for DAPI (cyan) and phalloidin (magenta). Scale bar = 10 μm. Right: Quantification of number of nuclear γH2AX.1-positive foci (N=1 experiment, n cells: Ctrl BMDC=88, CK666 D7-D10=153, CK666 D10 (2h)=108, Ctrl vs CK666 D7-D10 ****p<0.0001, Ctrl vs CK666 D10 (2h), ns p=0.1601, CK666 D7-D10 vs CK666 D10 (2h) ** p=0.0014; mean ± SD; ordinary one-way ANOVA, Tukey’s multiple comparison test). (B) Left: Representative confocal images of γH2Ax immunostaining in BMDCs at day 8 of culture. Scale bar = 10µm. Right: Quantification of nuclear mean γH2Ax signal in BMDCs at day 8 of culture (n ctrl= 65, n KO= 55, ****p<0.0001; mean ± SD; unpaired, two-sided t-test). (C) Flow cytometry plots of CD11c expression control and *Arpc4*KO BMDC cultured until day 7, 8, 9, 10 (n= 3 cultures ctrl, 3 cultures KO/ experiment/ experiment, ns p = 0.4857; mean ± SD; two-sided Mann-Whitney U-test). (D-H) RNA seq of BMDC from ctrl (black dots) and KO (magenta dots) after 7, 8, 9 or 10 days of culture. (D) PCA of highest variable genes (500), ellipse indicates similar condition. (E) GO Biological Processes (BP) enriched in differentially expressed (DE) genes, i.e. upregulated in KO samples compared to WT for each day. At day 10 there are no enriched pathways. (F) Heatmap of all DE genes with log2 fold change > 1. K-means clustering of genes. (G) Expression of each gene along time. Genes are grouped by cluster. (H) Hallmark pathways enriched in gene clusters 2 and 4. (I) Gene set enrichment analysis (GSEA) of a senescence gene signature (Aging Atlas Consortium 2021) was performed on bulk RNA-seq data across multiple time points (Days 7–10). Bubble plot showing normalized enrichment score (NES) for the AgingAtlas signature at each time point. Bubble size represents −log10 false discovery rate (FDR), and color indicates NES (red, positive enrichment; blue, negative enrichment on KO samples).

In line with this result, we found that day 10 control and *Arpc4*^KO^ DCs showed no to very little differences in gene expression, in contrast to cells at days 7-9 of differentiation (Fig. S5B). In particular, the enrichment of gene ontology (GO) term (biological process) analysis using the GO database revealed that day 8 and 9 *Arpc4*^KO^ cells displayed increased expression of immune-related biological processes, such as cytokine signaling and response to interferons (IFNs), as compared to their control counterpart (Fig. 5E). Similar results were obtained using Gene Set Enrichment Analysis (GSEA, Fig. S5C). K-means clustering of the genes differentially expressed (DEGs) between the two genotypes at the distinct developmental timepoints generated 7 different clusters (Fig. 5F,G; Fig. S5D). DEGs from clusters 2, 3, and 5 were elevated during the entire differentiation process in *Arpc4*-deficient cells (Fig. 5F-H; Fig. S5D). Cluster 2 included genes linked to IFN signaling and responses (Fig. 5F-H), suggesting that bone marrow cells might use Arpc4 to limit IFN signaling during DC differentiation. These results suggest that the events of nuclear envelope ruptures observed in *Arpc4*^KO^ DCs might lead to the production of type I IFNs through activation of the cGAS-STING pathway (Ablasser & Chang, 2019), a process associated with cellular aging (Gulen et al., 2023). However, we found that abrogation of STING function in *Arpc4*^KO^ mice did not rescue LC survival, excluding the possibility that aberrant STING activation triggers LC loss in these animals (Fig. S5E,F).

In addition to altered IFN and cytokine signaling, *Arpc4*^KO^ DCs also displayed significant differences in the expression levels of genes related to cell division. Indeed, cluster 4 included genes associated with mitotic progression (Fig. 5F-H), which were higher at days 7 and 8 of differentiation in *Arpc4*^KO^ DCs (Fig. 5G). Furthermore, GO Biological Process enrichment analysis highlighted an enrichment of mitosis- and cytokinesis-associated genes in day 7 *Arpc4*^KO^ DCs (Fig. 5H). GSEA of *Arpc4*^WT^ and *Arpc4*^KO^ DCs at days 7 to 10 of differentiation demonstrated the enrichment of a senescence-related signature (Aging Atlas Consortium, 2021) in *Arpc4*^KO^ cells (Fig. 5I), with consistently higher number of senescence-related genes being upregulated following ArpC4 loss (Fig. S5G). Together, these data open the possibility that bone-marrow precursors use Arp2/3 to limit IFN signaling as well as to progress through mitosis while differentiating into DCs, counteracting a permanent cell cycle arrest.

### Arpc4^KO^ DCs display altered expression of DNA repair factors, in particular during proliferation

Our results show on one hand that *Arpc4*^KO^ DCs exhibit aberrantly shaped and fragile nuclear envelopes, and on the other hand, that their transcriptional program during differentiation is characterized by increased IFN signaling and altered expression of genes related to cell division. We therefore hypothesized that in the absence of Arp2/3, differentiating bone-marrow precursors might encounter problems during cell division that lead to accumulation of DNA damage and aberrantly shaped nuclei. Such nuclei are prone to envelope ruptures, which in turn activate innate nucleic acid sensing pathways and downstream IFN production and signaling. In support of this hypothesis, it was recently shown in U2OS and HeLa cancer cells that Arp2/3 helps stabilizing replicative forks to allow the recruitment of DNA repair proteins during DNA replication, including RPA70 and ATR (Lamm et al., 2020; Palumbieri et al., 2023; Nieminuszczy et al., 2023). Accordingly, inhibition of the Arp2/3 complex in these cells attenuates replicative stress and accumulation of DNA damage. To investigate whether such mechanism could apply to *Arpc4*^KO^ DCs, we stained them with distinct markers for DNA double-strand breaks and for proteins of the DNA repair machinery. We observed that developing (day 8) *Arpc4*^KO^ DCs displayed increased staining for the 53BP1 DNA damage marker, in addition to γH2Ax (Fig. 6A,B). In contrast, staining for RPA70, which recruits the DNA repair kinase ATR to replicative forks, was found to be decreased in these cells (Fig. 6A,B). Accordingly, nuclear levels of ATR were also considerably diminished (Fig. 6A,B), opening the possibility of a defect in the recruitment of this kinase essential for DNA repair. These results strongly suggest that Arp2/3-dependent actin nucleation is needed to safeguard genome integrity in differentiating DCs.

**Figure 6:**
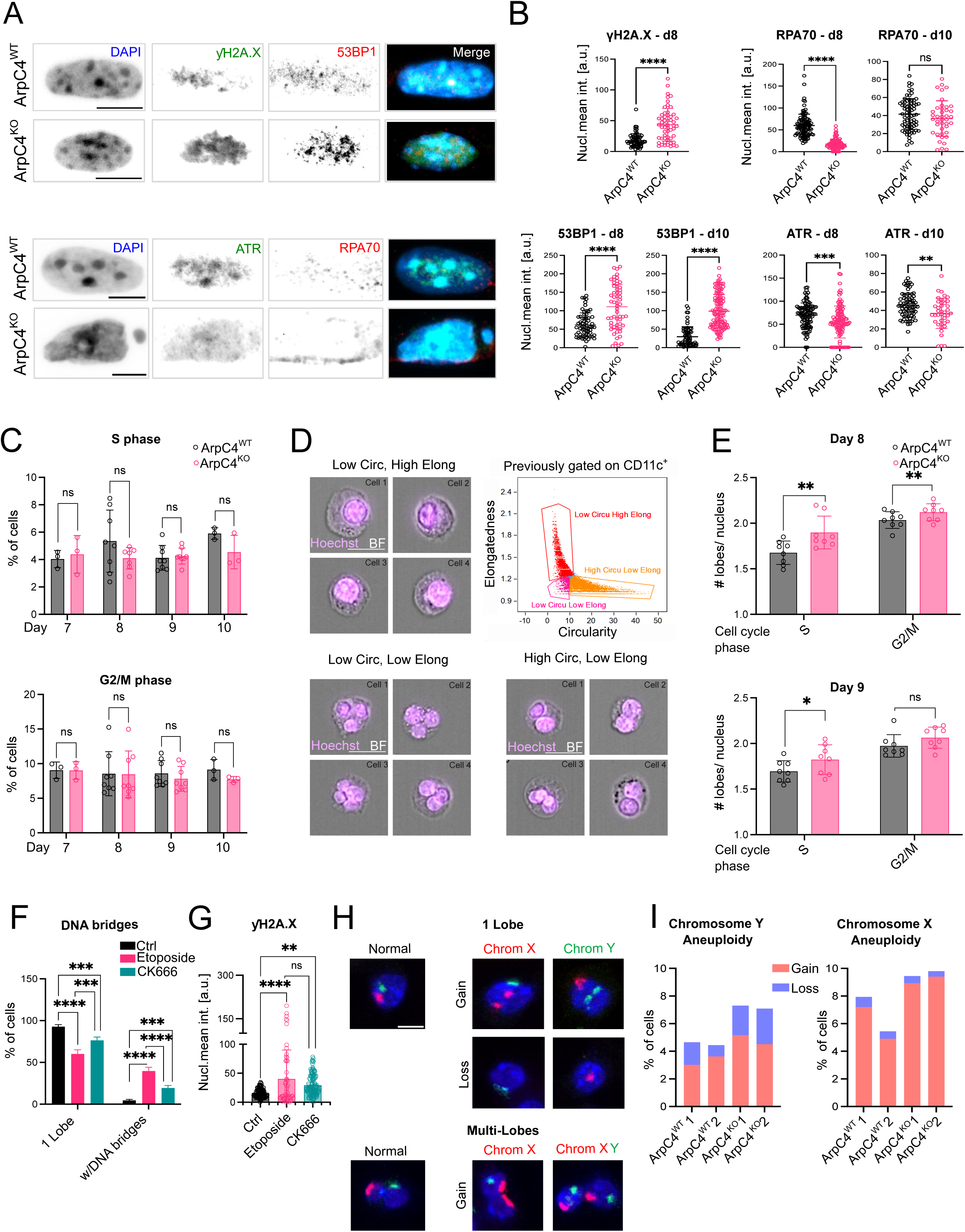
*Arpc4* deficiency leads to altered expression of DNA repair factors in BMDCs. (A) Representative confocal images of BMDC nuclei inside 8 μm × 5 μm fibronectin-coated µchannels. Upper panel: stained (from left to right) for DAPI, γH2Ax, 53BP1, and merge. Bottom panel: with DAPI, ATR, RPA70, and merge for both genotypes. Scale bars = 6 μm. (B) Quantification of nuclear mean fluorescence intensity of immunostainings for 53BP1 at day 8 of culture (n ctrl= 65, n KO= 58, ****p<0.0001; mean ± SD; unpaired, two-sided t-test), 53BP1 at day 10 of culture (n ctrl= 65, n KO= 124, ****p<0.0001; mean ± SD; unpaired, two-sided t-test), ATR at day 8 of culture (n ctrl= 103, n KO= 111, ***p=0.0005; mean ± SD; unpaired, two-sided t-test), ATR at day 10 of culture (n ctrl= 74, n KO= 41, **p=0.0022; SD; two-sided Mann-Whitney U-test); RPA70 in BMDCs at day 8 of culture (n ctrl= 101, n KO= 101, ****p<0.0001; mean ± SD; unpaired, two-sided t-test), RPA70 at day 10 of culture (n ctrl= 101, n KO= 115, ns; mean ± SD; unpaired, two-sided t-test). (C) Quantification of the percentage of BMDCs in S phase (upper panel) and G2/M (lower panel) at different days of culture. (D) Representative images of single BMDCs acquired on FACs-Streamer stained with Hoechst (magenta), overlay with bright field (BF). Upper left panel: Low circularity, high elongation, defines 1 Lobe population. Lower left panel: low circularity, low elongation, defines multi-lobes population. Lower right panel: High circularity, low elongation, defines 2-lobes population. Upper right panel: Gating strategy shows CD11c^+^ cells, three gates were used to define 1 lobe, 2 lobe or multi lobe cells according to their morphology values; elongation (Y axis) and circularity (X axis). Scale bar = 7 μm. (E) Quantification of the mean lobe number on BMDCs cultured until day 8 observed in S phase (n = 2 experiments/ BMDC cultures from 4 mice ctrl, 4 mice KO/ experiment, **p=0.0046; mean ± SD; unpaired, two-sided t-test). Mean lobe numbers of BMDCs cultured until day 8 observed in G2/M phase (**p=0.0084; mean ± SD; unpaired, two-sided t-test) for both genotypes (upper graph). Mean number of lobes of BMDCs cultured until day 9 observed in S Phase (*p=0.0412; mean ± SD; unpaired, two-sided t-test). Mean number of lobes of BMDCs cultured until day 8 observed in G2/M (ns; mean ± SD; unpaired, two-sided t-test) for both genotypes (lower graph). (F) Quantification of the percentage of cells with nuclei displaying 1 lobe or multi-lobes connected by DNA bridges per total cells analyzed in BMDCs from Ctrl mice, cells treated overnight with Etoposide and CK666 (N=3 independent BMDC cultures, total n ctrl=236, n Etoposide=142, n CK666=193. 1 Lobe: Ctrl. vs Etoposide ****p<0.0001, Ctrl vs. CK666 ***p=0.0004, Etoposide vs. CK666 ***p=0.0004. Multi-Lobes/with DNA bridges: Ctrl. vs Etoposide ****p<0.0001, Ctrl vs. CK666 **p=0.0008, Etoposide vs. CK666 ****p<0.0001; mean ± SD; unpaired, two-way ANOVA, Tukey’s multiple comparison test). (G) Quantification of nuclear mean fluorescence intensity of γH2Ax in BMDCs from Ctrl B6J mice treated overnight with Etoposide and CK666 (n ctrl= 129, n Etoposide= 54, n CK666= 177, Ctrl. vs Etoposide ****p<0.0001, Ctrl vs. CK666 **p=0.0066, Etoposide vs. CK666 ns p=0.1198; mean ± SD, ordinary one-way ANOVA). (H) Representative maximum intensity z-projection images of interphase nuclei of cells isolated from KO and ctrl mice. Chromosomes are stained using whole chromosome FISH probes (chromosome X in red and chromosome Y in green). Nuclei were stained with DAPI. Examples for euploid (left, normal) and aneuploid (right; gain and loss of chromosomal regions) nuclei with a single lobe are shown in the upper panels and multi-lobe nuclei are shown in the lower panels. Nuclei were stained with DAPI. Scale bar =5 μm. (I) Quantification of the percentage of aneuploid cells per total cells analyzed in each condition in two independent biological replicates showed in (H). Percentage of cells showing Y and X aneuploidy (N= independent cultures from 2 ctrl and 2 KO mice, total n WT1= 1096, WT2=1102, KO1=1355, KO2=776 cells, data were compared using logistic regression (GLM) with binomial distribution for Y; ***p= 0.000203 and for X; ***p=0.000541).

To evaluate whether these defects in DNA repair factors compromised the ability of *Arpc4*^KO^ cells to progress through the cell cycle during differentiation, we analyzed them in a FACSTREAMER, which combines flow cytometry with cell imaging. We found that only a small proportion of developing DCs (CD11c^+^) were in S (< 10%) and G2/M phases (5-15%) at any differentiation timepoint (Fig. 6C). These percentages were similar for control and *Arpc4*^KO^ DCs, suggesting that Arpc4 deficiency did not abrogate the cell cycle of developing DCs, consistent with *Arpc4*-mutant LCs being able to proliferate during postnatal skin development (Fig. 2). However, when analyzing the nuclear shapes of *Arpc4*^KO^ DCs, we observed that the percentage of cells with deformed nuclei was considerably enhanced in *Arpc4*^KO^ cells (Fig. 6D,E; Fig. S6A). This phenotype was spotted throughout S and G2/M phases but more pronounced in S phase, in agreement with this phase relying at most on the DNA repair machinery for genome replication. Hence, developing *Arpc4*^KO^ DCs that undergo proliferation display aberrant nuclear shapes more often than control cells. Together, these results suggest that although differentiating *Arpc4*^KO^ DCs start accumulating DNA damage upon proliferation, they succeed in progressing through the cell cycle. This is consistent with a model where Arp2/3 promotes DNA repair and nuclear integrity at least in part during the proliferation of mononuclear phagocytes *ex vivo* and *in vivo*.

In cancer cells, DNA damage resulting from defective repair can lead to altered nuclear architecture and envelope integrity (Kidiyoor et al., 2020), prompting us to investigate whether the aberrantly shaped nuclei of *Arpc4*^KO^ DCs could result from the DNA damage generated during proliferation. To address this question, we used etoposide to induce DNA damage during the differentiation of wild-type BM-derived DC cultures. This small molecule is known to induce DNA single- and double-strand breaks by inhibiting the topoisomerase II (Muslimovic et al., 2009). We found that etoposide treatment of differentiating DCs considerably increased the number of cells displaying multilobulated nuclei and DNA bridges (Fig. 6F). A similar effect was observed with the pharmacological Arp2/3 inhibitor CK666. As expected, both drugs increased DNA damage as shown by γH2Ax immunostaining (Fig. 6G). Therefore, treatment with either etoposide or CK666 phenocopy the nuclear shape alterations observed upon Arpc4 gene deletion, pointing to aberrantly shaped nuclei in Arpc4-deficient DCs being the consequence of elevated DNA damage. Overall, these results are consistent with a model where Arp2/3 promotes DNA repair in mononuclear phagocytes, helping these cells maintaining nuclear integrity upon proliferation. Accordingly, we further observed that *Arpc4*^KO^ DCs display increased Y chromosome aneuploidy (Fig. 6H,I), reinforcing our conclusion that Arp2/3 is needed to maintain genome integrity upon cell division in these cells. Of note, our data do not exclude that Arp2/3 could also contribute to the maintenance of genome integrity in non-dividing DCs.

### Arpc4 deficiency selectively compromises myeloid cell populations that rely on proliferation for tissue colonization

From this working model, it can be predicted that *in vivo* Arpc4 deficiency should preferentially affect the myeloid cell populations that, as LCs, rely on cell division to colonize or survive within their tissue of residency. In contrast, myeloid cell populations that proliferate prior to tissue colonization should remain unaffected in *Arpc4*^KO^ mice. To test the validity of these predictions, we first compared dermal cDC subsets in control and *Arpc4*^KO^ animals using flow cytometry. Indeed, these cells develop from preDCs, which acquire CD11c, and should thus delete the *Arpc4* gene prior to reaching the skin where they differentiate with at most one or two proliferation cycles (Cabeza-Cabrerizo et al., 2019). We found that the numbers of dermal cDC1s and cDC2s in unchallenged skin remained unchanged upon Arpc4 loss (Fig. 7A,B; Fig. S7A for gating strategy). To assess the functional consequence of ArpC4 loss in challenged skin *in vivo*, we performed skin painting with the contact sensitizer fluorescein isothiocyanate (FITC). Detection of cutaneous-derived phagocytes within draining lymph nodes revealed that migration of both *Arpc4*^KO^ DCs and LCs to lymph nodes was impaired (Fig. 7C-E), confirming a role of Arp2/3 for immune cell migration to lymphatic organs following chemical challenge. Similar differences were observed when analyzing steady-state migration of skin DCs, in agreement with our recent findings showing that Arp2/3 controls this process by activating the lipid metabolism enzyme cPLA2 (Alraies et al., 2024). Of note, loss of LCs was not observed in mice lacking cPLA2 in the CD11c-derived lineage (Fig. S7B), indicating that the role of Arp2/3 in their maintenance involves a distinct pathway.

**Figure 7:**
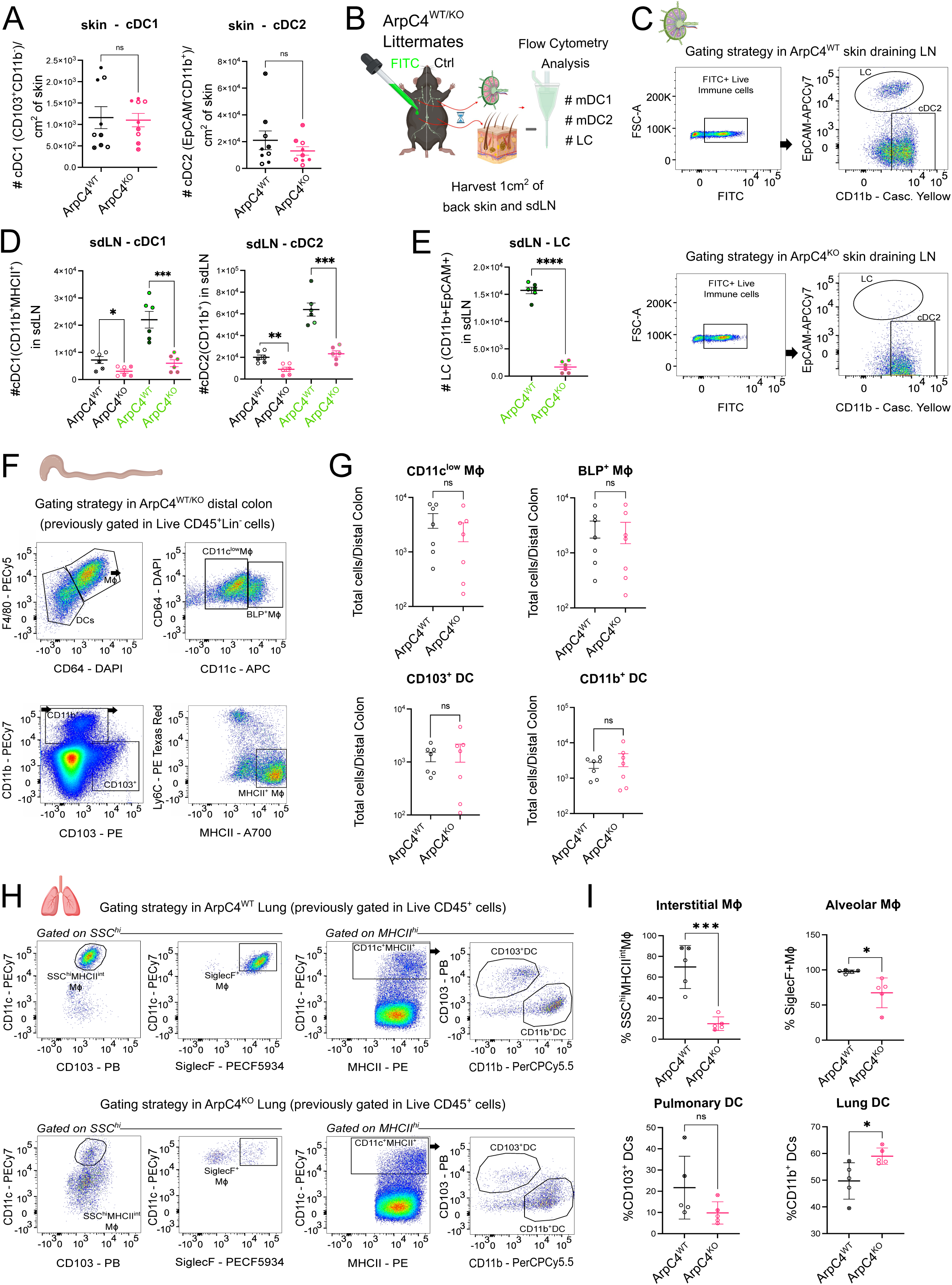
Lung alveolar macrophages, but not skin DCs and colon macrophages, rely on Arp2/3. (A) Quantification of cDC1 and cDC2 skin-resident dendritic cell compartments per cm^2^ of skin. cDC1: 3 experiments with 3 mice / genotype / experiment, n ctrl= 9, n KO= 9, ns, p= 0.8490; mean ± SD; unpaired, two-sided t-test); cDC2: 3 experiments with 3 mice / genotype / experiment, n ctrl= 9, n KO= 9, ns, p=0.3172; mean ± SD; unpaired, two-sided t-test). (B) Scheme of FITC-painting experiment. Skin and LN harvested from ctrl (untreated side) and FITC-treated samples (24 hours post-painting) were prepared for flow cytometry analysis. (C) Gating strategy on sdLN in FITC experiments: first CD45^+^ immune cells were identified, and then only FITC^+^ cells were considered for the rest of the analysis done exactly like in Ctrl cells. (D) Quantification of DC migration to LNs. Ctrl (untreated) side is shown on the left, whereas on the right, green letters indicate samples from FITC-treated side (2 experiments with 3 mice / genotype / experiment, n ctrl= 6, n KO= 6). cDC1, ctrl side; *p=0.0197; mean ± SD; unpaired, two-sided t-test, FITC-treated side; ***p=0.0007; mean ± SD; unpaired, two-sided t-test. cDC2, ctrl side; **p=0.0032; mean ± SD; unpaired, two-sided t-test, FITC-treated side; ***p=0.0001; mean ± SD; unpaired, two-sided t-test. (E) Quantification of LC migration from FITC-treated skin to sdLN (2 experiments with 3 mice / genotype / experiment, n ctrl= 6, n KO= 6, ****p<0.0001; mean ± SD; unpaired, two-sided t-test). (F) Gating strategy in distal colon at steady state for *ArpC4*^WT^ and *ArpC4*^KO^ (only WT is shown), CD11b^+^ cells were used to identify MHCII^+^ MΦ, which were subsequently separated on colonic (monocyte-derived) F4/80^+^CD64^+^ MΦs and F4/80^−^CD64^−^ DCs; this last one was then separated in CD11c^low^ and CD11c^high^ MΦs. (G) Quantification of colonic (monocyte-derived) ctrl vs. *Arpc4*^KO^ macrophages (2 experiments with 3-4 mice / genotype / experiment), CD11c^low^ MΦs (ns p = 0.3829; mean ± SD; two-sided Mann-Whitney U-test), CD11c^high^ MΦs (ns p = 0.8048; mean ± SD; two-sided Mann-Whitney U-test), CD103^+^DCs (ns p = 0.9720; mean ± SD; two-sided Mann-Whitney U-test) and CD11b^+^DCs (ns p = 0.9755; mean ± SD; two-sided Mann-Whitney U-test). (H) Gating strategy in lung at steady state for *ArpC4*^WT^ (upper panel) and *ArpC4*^KO^ (lower panel). MΦs with high granularity (SSC^hi^) and intermediate MHCII signal were identified (SSC^hi^ MHCII^int^MΦ, referred to as interstitial MΦs) as well as Siglec F^+^ MΦ (alveolar MΦs). MHCII^hi^ gate was used to define pulmonary DCs (CD103^+^DCs) and lung DCs (CD11b^+^DCs). (I) Quantification of lung macrophages shown in (H). Interstitial MΦs (***p = 0.0005; mean ± SD; unpaired, two-sided t-test), alveolar MΦs (*p = 0.0133; mean ± SD; unpaired, two-sided t-test), pulmonary DCs (ns p = 0.1278; mean ± SD; unpaired, two-sided t-test) and lung DCs (*p = 0.0239; mean ± SD; unpaired, two-sided t-test). Organ schemes in B, C, F, and H used from Biorender (www.biorender.com).

These results emphasize that while LCs and DCs use Arp2/3 for efficient migration to lymph nodes, this complex is dispensable for the formation of cDC1 and cDC2 pools from preDCs in the mouse skin. In line with these results, we further observed that the numbers of colonic macrophages were not affected in *Arpc4*^KO^ mice (Fig. 7F,G; Fig. S7C for genotyping of sorted colon macrophages verifying successful deletion of ArpC4 in these cells; Fig. S7D for gating strategy), consistent with these cells differentiating from monocytes without involving extensive cell proliferation (Guilliams et al., 2014; Bain et al., 2022). In sharp contrast, when analyzing lung alveolar macrophages, which proliferate to self-maintain during adult life (Imperatore et al., 2017; Bain et al., 2022), we found that they were strongly decreased in numbers in *Arpc4*-deficient mice (Fig. 7H,I; Fig. S7E for gating strategy). Together, these data strongly suggest that Arp2/3 is preferentially required for the maintenance of the pool of myeloid cells that rely on cell division to colonize or to self-maintain within their tissue of residency, and that in the absence of this complex, these cells die prematurely.

## DISCUSSION

We here show that genetic deletion of Arpc4 leads to the loss of LCs in the skin of both the ear and tail of young adult mice. This is associated with accumulation of DNA damage in these cells, which further exhibit aberrantly shaped nuclei and, consistently, decreased expression of the nucleoskeleton proteins Lamin A/C and B1. DCs derived *in vitro* from the bone-marrow recapitulate the defects observed in *Arpc4*^KO^ LCs, allowing us to investigate underlying mechanisms. Our results suggest that Arp2/3 plays a role in the maintenance of both DNA and nuclear envelope integrity and concomitant nuclear shape, which if altered can lead to premature cellular aging.

Nuclear actin networks have been reported for many years and Arp2/3 has recently been implicated in the mobility of double-strand DNA breaks to the nuclear periphery for productive repair (Schrank et al., 2018; Lamm et al., 2020). However, most of these studies were restricted to cultured cells. The findings made in the present study suggest that the recently reported role of branched actin networks and Arp2/3 in DNA repair might indeed play a key physiological role in maintaining the genome integrity of defined myeloid cell populations *in vivo*. Importantly, our results using DCs derived from the bone-marrow suggest that this role of Arp2/3 might particularly apply to DCs that proliferate in the course of maturation and differentiation. This is further supported by our findings of a role for the Arp2/3 complex in the homeostasis of myeloid cells that rely on proliferation for tissue colonization or residency (i.e., LCs and alveolar macrophages), in contrast to post-mitotic immune cell populations (i.e., colonic macrophages and skin cDC1 and cDC2). Of note, we cannot exclude that ontogeny might also contribute to the differences observed between skin conventional DCs and bone-marrow-derived DCs, as we used for this study a differentiation protocol that relies on GMCSF and most likely leads to DCs differentiation from monocyte precursors rather than from DC ones. GMCSF-dependent bone-marrow-derived DCs are thus closer to cells from the macrophage lineage, which could further explain their phenotypic similarities with LCs.

Our data are fully consistent with DNA replication having a strong dependency on DNA repair capacities. Noticeably, Arp2/3 has been involved in the opening of replication forks for recruitment of the DNA repair machinery during replication (Lamm et al., 2020; Palumbieri et al., 2023; Nieminuszczy et al., 2023), and its inhibitor Arpin was recently shown to inhibit DNA homology repair (Simanov et al., 2024). Surprisingly, we found that both *Arpc4*-deficient DCs and LCs can nonetheless progress through the cell cycle, suggesting that DNA damage accumulation upon division in these cells is not sufficient to fully activate cell cycle checkpoints. Interestingly, it was recently shown that Formins, another class of actin nucleators, can also be found in the nucleus where they modify nuclear actin networks (Parisis e al., 2017; Ulferts et al., 2024; Ulferts & Grosse, 2024), opening the possibility of a local compensation for the loss of Arp2/3.

It seems well possible that the defects observed upon *Arpc4* loss engage a bidirectional link between DNA damage and decreased nuclear envelope integrity, as suggested by our result that induction of DNA damage with etoposide during DC differentiation led to aberrantly shaped nuclei. In cancer cells, replication stress or defective DNA repair can result in altered nuclear architecture, reduced nuclear integrity and blebbing nuclei (Lamm et al., 2020; Kidiyoor et al., 2020; Eskndir et al., 2024). A recent study further reported that DNA damage-induced nuclear changes result in more fragile nuclear envelopes that easily break under mechanical constraints (Kovacs et al., 2023). On the other hand, we and others have shown that nuclear envelope deformations and rupture trigger the accumulation of DNA damage due to the exposure of nuclear DNA to cytoplasmic nucleases (Raab et al. 2016; Denais et al., 2016; Shah et al., 2021), e.g. by driving Trex1-dependent DNA damage as shown in human breast cancer cells (Nader et al., 2021). Although the precise mechanisms underlying Arp2/3-dependent nuclear envelope and genome integrity remain open, we found that the deformed nuclei in *Arpc4*^KO^ cells are more fragile and prone to envelope rupture even in the absence of external physical constraints. We suspect that accumulation of DNA damage upon division of *Arpc4*^KO^ cells could be further amplified by exposure to these enzymes. This hypothesis is in good agreement with our RNAseq results showing enhanced type I IFN signaling in BMDCs lacking *Arpc4*, in addition to altered cell division signatures. Based on all these findings, we propose a model where proliferating *Arpc4*^KO^ LCs and DCs start accumulating discrete events of DNA damage because of impaired DNA repair during cell division. This could result from compromised fork opening, as previously described in cell lines. However, although these DNA damage events would not prevent progression through the cell cycle, they would result in nuclear deformations, as observed in LC networks of *Arpc4*^KO^ mice *in vivo* and in *Arpc4*^KO^ DCs *in vitro,* and possibly mitotic errors that have previously been associated with nuclear deformation (Hwang et al., 2019). Such deformed nuclei would experience frequent events of nuclear envelope rupture, entering a DNA damage amplification loop. This model provides a putative explanation for *Arpc4*^KO^ LCs successfully dividing upon epidermal colonization at birth but accumulating DNA damage only four weeks later, followed by a progressive decline of LCs.

Importantly, we do not exclude that effects of Arp2/3 other than maintenance of genome integrity could contribute to the loss of LCs and alveolar macrophages *in vivo*. This is particularly relevant for LCs, as we found that they exhibited decreased dendritic branching when lacking *Arpc4*. The branched morphology of LCs was also shown to depend on the small GTPase Rac1, a well-known activator of Arp2/3, and is potentially needed for proper interaction of LCs with surrounding keratinocytes during tissue colonization (Park et al., 2021). Similarly, Cdc42, another small GTPase that turns on Arp2/3 activity, was found to be necessary for maintenance of LC numbers in the mouse skin (Luckashenak et al., 2013). These results point towards Arp2/3 being required not only for genome integrity in LCs, but also for nucleation of peripheral actin networks needed for these cells to interact with their cellular environment. Interestingly, we have shown that there is a competition for Arp2/3 subcellular pools between the cell periphery and the perinuclear area where the centrosome is located (Obino et al. 2016; Inoué et al. 2019). Such competition is promoted by cell adhesion to the environment, which leads to Arp2/3 recruitment at the cell periphery and results in its depletion from the perinuclear space. In lymphocytes, this facilitates centrosome detachment from the nucleus and polarization towards the immune synapse (Obino et al. 2016; Inoué et al. 2019). Whether such competition is restricted to cytoplasmic Arp2/3 or could involve the pool of intranuclear actin in LCs to help coordinate cell-cell adhesion and genome integrity in time and space is an intriguing hypothesis that may deserve further investigation. Interestingly, recent work performed in motile T lymphocytes or DCs highlights an actin regulatory loop between the central actin and leading edge actin as well as a role of the Guanine Exchange Factor DOCK8 in formation of a perinuclear actin pool and maintenance of nuclear integrity during migration in confinement (Shen et al., 2024; Reis-Rodrigues et al., 2024). Our data together with recent reports opens the possibility that Arp2/3 complex members serve various roles in different immune cell contexts *in vivo*. While chemical inhibition of the Arp2/3 complex caused cell death in crowding neutrophils (Glaser et al., 2023), Arp2/3 loss in mast cells resulted in cell death that involved integrin-mediated mechanocoupling to the underlying extracellular matrix (Kaltenbach et al., 2024). Similarly, the development and dynamics of microglial cells has recently been shown to engage Arp2/3 function (Scott-Solache et al., 2026; Safaiyan et al., 2026; Paulson et al., 2026). A recent study that employed deletion of ArpC5 and ArpC5L in the murine hematopoietic lineage revealed a requirement of ArpC5 for phagocytosis and killing of intracellular bacteria, resulting in intestinal inflammation (Vasconcellos et al., 2025). While that study did not focus on subcellular ArpC5 pools, we observed that a subfraction of BMDCs shows Arpc5-positive foci within their nuclei (Fig. S3F,G), opening the possibility that Arp2/3 components may in principle act in the nucleus in these cells.

At present, the exact cellular mechanism of how *Arpc4*^KO^ LCs are progressively lost from the epidermis remains open. A previous study reported the dependency of adult LCs on the late endosomal adaptor molecule p14 (LAMTOR2). Upon LAMTOR2 deficiency, the initial burst of proliferation is followed by reduced proliferation and increased apoptosis (Sparber et al., 2014). While we detected a temporary trend for increased apoptosis of LCs in ear and tail epidermis at the age of 4 weeks, this trend disappeared at the age of 6 weeks. We assume that only a short time passes until LCs with impaired genome integrity will be cleared from the system, making it difficult to detect γH2Ax-positive cells that are positive for markers of cell death. Unfortunately, because bone-marrow-derived cultures of DCs have a short lifespan, unlike LCs *in vivo*, these cell death mechanisms could not be investigated in detail in the present study. However, we also note that no robust apoptotic signatures were detected in *Arpc4*^KO^ DCs *in vitro* (Fig. S5H). We can thus not exclude that damaged LCs are eliminated by cell death pathways other than apoptosis.

Remarkably, the phenotypes of *Arpc4*^KO^ LCs and DCs share many similarities with the phenotype of cells that age prematurely, including altered nuclear shapes, fragmentation of the Lamin nucleoskeleton, nuclear envelope rupture, enhanced IFN signaling, loss of genome integrity and aneuploidy. Accordingly, Arp2/3 deficiency in cell lines was shown to trigger a cell senescence transcriptomic signature (Wu et al., 2013). This is in line with our observation of an enriched senescence signature found in *Arpc4*^KO^ DCs as well as an enhanced E2F pathway, which is associated with cell senescence. These results are consistent with our findings pointing to LCs and bone-marrow-derived DCs from *Arpc4*-deficient animals exhibiting an accelerated cellular aging phenotype. Of note, whether Arp2/3 activity declines in aging cells and tissues and thereby contributes to accumulation of DNA damage is not addressed in the present study. Collectively, our results provide insights into Arp2/3-dependent control of LC maintenance and genome integrity and highlight an important *in vivo* role for actin regulation in proliferating tissue-resident myeloid populations.

## MATERIALS AND METHODS

### Mice

An *Arpc4* conditional allele was used as described in Rivera et al. 2022, and mice with deletion of ArpC4 in CD11c-derived lineages (*CD11c*Cre^wt/+^;*Arpc4*^flox/flox^ (Itgax=CD11c) and cPLA2 mutant animals (*CD11c*-Cre;*Pla2g4a*^flox/flox^) were generated as described in Alraies et al, 2024. STING-GT (Goldenticket; Sauer et al., 2011) were crossed to *CD11c*Cre^wt/+^;*Arpc4*^flox/flox^ mice. Genetic background stability was ensured by retro-crossing the line with C57BL6/J mice, obtained from Charles River (000664). Animals were bred and maintained in the animal facilities at Institut Curie, Paris, France and Saarland University, Homburg, Germany, for the indicated experiments. Mice were handled according to the European and French National Regulation for the Protection of Vertebrate Animals used for Experimental and other Scientific Purposes (Directive 2010/63; French Decree 2013-118, Authorization DAP number 43530-2023051216135493 v2 given by French Authority), and according to federal guidelines in Germany. FACS experiments were performed using 8- to 14-week-old female mice, in the case of chromosome x and y aneuploidy analysis only male mice were used. Littermates or age-matched mice were used as controls for all experiments involving knockout animals. Mice were maintained under specific pathogen-free (SPF) conditions at Institut Curie animal with free access to food, water and housed in a 12-h light/12-h dark environment and (MUCEDOLA, 4RF25SV Aliment Pellets 8 × 16 mm irradiated 2.5 Mrad and DietGel Recovery, 2 oz/DietGel Boost, 2 oz). In Homburg, all mice were housed and maintained according to federal guidelines within the SPF animal facility in Building 61.4 of the Saarland University under an approved breeding license (§11 Az. 2.4.1.1-Iden). All experiments were performed in line with German animal welfare laws as well as institutional guidelines and were reported to the responsible authorities.

### DNA isolation

Tails of P7 animals were collected after euthanasia. Following, an epidermal split was performed as described for tail whole mounts which allowed to only isolate DNA from the LC containing epidermis. Subsequently, samples were lysed in tissue lysis buffer (100 mM Tris pH 8.0; 5 mM EDTA; 0.2% SDS; 200 mM NaCl) with freshly added proteinase K (5 µg/mL; Invitrogen) at 55°C for 30 min on a rocking station at full speed. After a vigorous vortex step samples were centrifuged at full speed for 10 min allowing the sedimentation of debris. The supernatant was mixed with same parts isopropanol for DNA precipitation and centrifuged again. The resulting pellet war washed in 70% ethanol and after a final centrifugation step resuspended in 200 µl TE-Buffer and gently shaken at 55°C for 30 min to dissolve the DNA.

### Genotyping

To test for the presence of the transgenes or recombination events of interest, PCRs using gene-specific custom-designed primers (Table S7) were performed with the ReadyMIX REDTaq (Sigma-Aldrich).

### Immunohistochemistry of tail epidermal wholemounts

Preparation and immunostaining of mouse tail wholemounts was performed as previously described (Braun et al., 2003). Briefly, mouse tail skin was peeled off the bone and incubated in 5 mM EDTA/PBS for 1.5 h at room temperature (RT) with gentle shaking at 37°C. After replacing half of the EDTA/PBS volume, the tissue was incubated for another 1.5 h. For tissue preparation of younger mice, incubation times were adapted. Next, the epidermis was separated from the dermis, leaving hair follicles attached to the epidermal sheet. Depilation was performed to eliminate autofluorescence caused by hair. The epidermis was fixed in pre-cooled acetone for 20 min on ice. For blocking, samples were incubated in PB buffer [0.5% skim milk powder, 0.25% fish skin gelatin (Sigma-Aldrich, cat. #9000-70-8), 0.5% Triton X-100 in HBS (20 mM HEPES pH 7.2, 0.9% NaCl)] or 5% BSA/PBS for 1 h at RT, followed by incubation with primary antibodies diluted in blocking buffer overnight at 4°C. The next day, samples were washed three times in 0.05% Triton X-100/PBS for 30 min. Samples were then incubated with secondary antibodies (diluted in blocking buffer) for 4 h at RT. Finally, washing steps for 30 min were repeated at RT and samples were mounted in Fluoromount G with the basal side facing upwards. Micrographs were acquired next to the dorsal midline. For details of antibodies used see Table S6.

### Immunohistochemistry of ear epidermal wholemounts

Preparation and staining of mouse ear epidermis wholemounts was performed as previously described (Ross et al., 1998). Briefly, mouse ears were depilated using Veet Hair Removal Cream and residual ventral cartilage was removed. Ears were split while floating on PBS. To separate dermis and epidermis, the skin was placed on 0.5 M ammonium thiocyanate for 25 min at 37°C or 5 mM EDTA/PBS for 1 h at 37°C with the dermal side down. For tissue preparation of younger mice incubation times were adapted. After removing the dermis from the epidermis using forceps, the epidermis was spread on the bottom of a 12-well plate, covered with ice-cold acetone, and fixed on ice for 20 min. Next, ear tissues were blocked in PB buffer or 5% BSA/PBS for 1 h at RT and subsequently incubated with primary antibodies overnight at 4°C. The next day, ear epidermis was washed three times in 0.05% Triton-X-100/PBS for 20 min at RT, followed by incubation with secondary antibodies (diluted in blocking buffer) for 2 h at RT. Finally, the ear tissues were washed three times in washing buffer at RT and then mounted in Fluoromount-G, with the basal side facing upwards. Per experiment only inner or outer parts of the ear were compared. Micrographs were acquired in central regions of the ears. For details of antibodies used see Table S6A,B.

### Immunocytochemistry of BMDCs

BMDCs were washed with PBS and incubated with 4% paraformaldehyde (PFA, Pierce, cat. #28906) diluted in PBS for 30 min at RT. After fixation, samples were washed and incubated with the primary antibodies (Table S5A) overnight at 4°C. After washing with PBS, cells were incubated for 2 h at 4°C with secondary antibodies respectively (Table S5B). For STED microscopy a special set of secondary antibodies was used (Table S5C). Finally, samples were washed three times with PBS and mounted with Fluoromount-G solution (Invitrogen, cat. #00-4958-02).

### Immunofluorescence microscopy - Tissues

Confocal images of epidermal sheets were acquired using a Zeiss LSM880 confocal laser scanning microscope and ZEN 2.3 SP1 software using a PL Apo 40×/1.3 Oil DIC UV-IR M27 or PL Apo 63x/1.4 Oil DICII objective. To analyze intranuclear signals, images were acquired using the Zeiss Airyscan detector. Images were acquired using identical imaging and background subtraction settings for both WT and KO samples for each marker used in the immunofluorescence experiments.

### Immunofluorescence microscopy - BMDCs

Images of immunostained BMDCs were acquired with a confocal microscope (Leica DMi8, SP8 scanning head unit) with a ×40/1.3-NA oil objective and a resolution of 1,024 × 1,024 pixels. Super-resolution images were acquired on an inverted STED microscope (Stedycon, Abberior Instruments, Göttingen, Germany) with a Leica 100X/1.4 Oil Plan Apo CS2 objective and a toroidal (“donut”) depletion pattern of the STED focus with a 775 nm laser. Images were deconvolved using theoretical point spread functions on Huygens Professional (Scientific Volume Imaging, The Netherlands, http://svi.nl). Nuclear images showing DNA bridges were acquired with a 3D SIM in a SR-SIM microscopy system. For this a Delta Vision OMX v4 microscope equipped with an Olympus 100X/1.4 Oil UPlan Apo objective and 3 EMCCD cameras was used. Image reconstruction was performed using SoftWoRx. ArpC5 subcellular localization was analysed using an Elyra 7 Structured Illumination microscope (Carl Zeiss, Germany) with 63x/1,46 Plan-Apochromat Oil objective and 2 cameras Edge 4.2 CLHS (PCO). In all cases, images were acquired using identical imaging and background subtraction settings for both WT and KO samples for each marker used in the immunofluorescence experiments.

### Quantitative analysis of LC densities in epidermal sheets

For analysis of LC densities in tail and ear epidermis of P1, P7, 4-weeks-old and adult mice, MHCII^+^ or CD207/langerin^+^ cells were counted and cell numbers per mm^2^ tissue was calculated.

### Voronoi diagrams

Voronoi diagrams were generated by using the CD207^+^ signal to produce a thresholded mask of LCs with an Image J customized macro (v2.9.0/1.53t, National Institutes of Health). Subsequently, the Voronoi tool was used to produce the diagrams.

### Morphological analysis of LCs and their nuclei *in vivo*

Cell profiler software (v.4.2.5, Broad Institute) was used to create masks and skeletons of individual LCs from maximum intensity projections of image acquisitions at 40x magnification. The skeletonized images were further analyzed using “analyze skeleton” plugin in ImageJ (v.1.54f, National Institutes of Health). Nuclei of single LCs (CD207^+^ cells) were classified using super-resolution images acquired with a Zeiss LSM880 with Airyscan. Because LCs exhibit elongated, nearly rectangular nuclei under homeostatic conditions, standard shape descriptors were not applicable. Therefore, each nucleus was evaluated individually and categorized as either normal or deformed.

### Analysis of apoptosis and proliferation in epidermal sheets

For analysis of apoptotic or proliferating LCs, partly depilated and acetone-fixed tail or ear epidermal whole mounts were blocked in PB buffer for 1 h at RT followed by incubation with primary antibodies diluted in PB buffer overnight at 4°C. The next day, samples were washed three times for 10 min each with 0.05% Triton X-100/PBS. Subsequently, samples were incubated with secondary antibodies (diluted in PB buffer) for 2 h at RT, followed by three washing steps (10 min each, RT) and mounting of samples in Fluoromount G, with the basal side facing upwards. Apoptotic cells in the remaining hair follicles served as internal positive control for cleaved caspase3 immunostaining, as apoptosis is a natural process occurring during catagen. Proliferating cells in the basal layer and remaining hair follicles served as internal positive control for Ki67 immunostaining, as these regions harbor proliferative active stem cell populations. For details of antibodies used see Table S6A,B.

### Analysis of DNA damage in epidermal sheets

For quantification of γH2AX-positive cells in ear and tail epidermis, maximum intensity projections of z-stack images of γH2AX, CD207 and DAPI stained whole mounts were used. γH2Ax-positive LCs (CD207^+^) were manually counted. Data from five individual animals per genotype were pooled to calculate the percentage of positive compartments.

### Flow cytometry analysis of DC subsets *in vivo* - Skin and skin-draining lymph nodes

DC subpopulations in skin-draining lymph nodes (LNs) were analyzed as described in Alraies et al, 2023. Briefly, 1 cm^2^ of back skin from 8-14 weeks old female wild-type and knockout littermates was dissociated in 0.25 mg/mL Liberase (Sigma, cat. #5401020001) and 0.5 mg/mL DNase (Sigma, cat. #10104159001) in 1 mL of RPMI (Sigma) and mechanically disaggregated in Eppendorf tubes, followed by incubation for 2 h at 37 °C and three-times filtration using a 100 µm cell strainer. Thereafter, blocking was performed in PBS supplemented with 0.5 % bovine serum albumin and 2 mM EDTA at 4 °C, followed by antibody labeling of cells in single cell suspension. In parallel, cells from inguinal and axial LNs were collected by mechanical disaggregation in RPMI medium supplemented with 0.5 mg/mL DNase and 1 mg/mL collagenase (Sigma, cat. #C2139) for 20 min at 37 °C. Single cells from LNs and skin were resuspended for blocking in PBS/BSA (see above), filtered three times using a 100µm cell strainer, blocked with Purified Rat Anti-Mouse CD16/CD32 (Mouse BD Fc Block, BD Biosciences, cat. #553142) and stained using the antibody mix listed in Table S1. Additionally, for the skin, an immune cell marker was added (Table S1). Subsequently, cells were filtered through 35 µm filter caps into 12 x 75 mm flow cytometry tubes (Dutcher, cat. #062985). Each sample was resuspended in 300 µL of PBS-0.5% bovine serum albumin (BSA)-1mM EDTA with 1 µL of D 4’,6-Diamidino-2-Phenylindole, Dilactate (DAPI, Biolegend, cat. #422801) (0.1 µg/mL) and 10 µL counting beads (Biolegend, cat. #424902), which were used to normalize the number of cells per sample. Flow cytometry was performed acquiring 0.5*10^6 live cells (DAPI-negative) per sample.

### Flow cytometry analysis of DC subsets *in vivo* - Lung

Lungs were dissected from 12-14 weeks old female wild-type and knockout littermates and stored in RPMI. Subsequently, lung tissues were mechanically disaggregated in RPMI supplemented with 0.4 mg/mL collagenase, 20 μg/mL DNAse I for 30 min at 37 °C. Afterwards, tissue mix was homogenized using a 100μm filter using PBS supplemented with 1% BSA and 1 mM EDTA and a 5mL syringe plunger. Red blood cells were lysed with red blood cell lysis buffer (Ozyme, cat. #42031) for five min at 4°C and reaction was stopped by adding heat-inactivated fetal bovine serum (FBS). Subsequently, cells were washed and filtered again through a 100 μm filter. The resulting single-cell lung suspension was stained with a 1:500 Live/Dead probe (APC-Cy7, Live/Dead, Invitrogen, cat. #L34992) diluted in PBS, blocked with Purified Rat Anti-Mouse CD16/CD32 and stained for flow cytometry analysis using the antibody mix listed in Table S2. 2*10^6 live cells per genotype were acquired and total cell counts were determined by addition of counting beads.

### Flow cytometry analysis of DC subsets *in vivo* - Distal colon

The analysis of distal colon cells was performed as described previously by Chikina et al, 2020. Briefly, colons were dissected and incubated on a magnetic stirrer in complete medium (CM, 2% heat-inactivated FBS in Ca2+, Mg2+-free Phenol Red 10X HBSS) with 1 mM DTT and EDTA at 37°C for 30 min. Subsequently, samples were incubated with 1 mM EDTA in 5% FBS/ PBS at 37°C for 10 min, followed by incubation with 15 mM HEPES in 1% FBS/PBS at RT for 7 min without agitation. Isolated tissues were collected and digested using 0.15 mg/mL Liberase and 0.1 mg/mL DNase I in HBSS at 37°C for 45 min with magnetic agitation. Single-cell suspensions were stained with Live/Dead probe diluted in PBS (LIVE/DEAD™ Fixable Aqua Dead Cell Stain Kit, ThermoFisher, cat. #L34965), blocked with Purified Rat Anti-Mouse CD16/CD32 and subsequently stained with antibodies listed in Table S3. Total cell counts were determined by addition to counting beads.

All samples were acquired on a LSRII cytometer (BD Biosciences) and processed with FlowJo version 10 software. Percentage and number of cells were plotted using GraphPad Prism version 9.

### Isolation and genotyping of CD11c+ gut macrophages

Colons were dissected from 11-weeks-old wild-type and knock-out littermates, as well as C57BL6/J mice and stored in 5% FCS-PBS on ice. They were then cut into 2 cm pieces and washed twice in 10 mL of DPBS. The epithelium was stripped out by incubating colon pieces in 15 mM HEPES (#15630-056 Gibco), 5 mM EDTA (#15575-038 Invitrogen), 5% FCS, 1 mM DTT (#D7997-1G Sigma), 1x HBSS (#H4385-100 Sigma) 30 min at 37°C under magnetic agitation at 600 rpm. Colon pieces were retrieved and washed twice with 10 mL of DPBS then incubated 10 min at 37°C under magnetic agitation at 600 rpm in 1 mM EDTA, 5% FCS, PBS (Wash A, 20 mL). Colon pieces were retrieved and washed twice with 10 mL of DPBS then incubated 7 min at RT in 15 mM HEPES, 1%FCS, PBS (Wash B, 10 mL). Again, two DBPS washes were realized, and remaining pieces were digested to extract immune cells from lamina propria by incubating in 2.5 mg Liberase, 0.5 mg DNase, 1x HBSS for 45 min at 37°C under magnetic agitation at 600 rpm. 20 mL of Wash A and of 1x HBSS were added on top, and the solution was filtered using a 100 µm pore filter and centrifuged for 7 min at 400g at 4°C. The supernatant was discarded, and cells were plated in a 96well-plate in FACS buffer (2 mM EDTA, 0.5% FCS, 0.5% BSA in DPBS). Cells were pelleted by centrifugation (320 x g, 5 min at 4°C) and then resuspended in Fc-block (1:100, #553142 BDBioscience) in FACS buffer for 15 min in the dark on ice. Cells were centrifuged again as before and resuspended in 100 µL of the antibody mix (Table S3) and then incubated on ice in the dark for 30 min. Then, 200 µL of FACS buffer were added, followed by centrifugation (320 x g, 5 min at 4°C). The supernatant was discarded, and an additional wash was performed. Cells were then resuspended in 100 µL of FACS buffer and DRAQ7 (1:1000, #D15106 ThermoFisher) was added right before sorting the cells. CD45.2+ CD64+ MHCII+ F4/80+ CD11c+ cells were sorted (1.105 cells for each sample) and DNA was extracted using the NucleoSpin tissue PureLinkTM kit (#K182001 Invitrogen). cDNA corresponding to the ARPC4 deleted band after Cre recombination was amplified (Table S7 for primers and cycling protocol) and the resulting products were loaded on 2% agarose gel. The intensity of the ArpC4 deleted band (147pb) was measured and normalized over the C57BL6/J (control) band.

### FITC painting to track DC and LC migration

Contact hypersensitivity was induced on the abdominal skin of isoflurane-anesthetized mice. The skin was shaved with a shaving machine. Thereafter, a 100 μL drop of 0.5% fluorescein isothiocyanate (FITC, Sigma Aldrich, cat. #3326-32-7), diluted in the sensitizing agent dibutyl phthalate (DBP, Sigma Aldrich, cat. #84-74-2) diluted 1:1 in acetone (Sigma Aldrich, cat. #67-64-1) was applied. Mice were analysed 24 hours following FITC treatment.

### BMDC extraction, culture maturation and transfection Bone marrow extraction

Bone marrow (BM) was isolated from tibias and femurs of 8- to 10-week-old male and female mice. Therefore, the epiphysis was cut off and bones were flushed with a syringe containing “DC medium” composed of IMDM medium (Sigma-Aldrich, cat#12440053) supplemented with decomplemented and filtered 10% FBS (Biowest), 20 mM/L glutamine (Gibco, cat. #25030081), 100 U/mL penicillin– streptomycin (Gibco, cat. #15070063), 50 μM 2-mercaptoethanol (Gibco, cat. #31350010) and 50 ng/mL of supernatant containing granulocyte–macrophage colony-stimulating factor (GM-CSF). GM-CSF was generated by transfected J558 cells and tested by enzyme-linked immunosorbent assay. Cell suspensions were centrifuged at 1300 rpm for 5 min. The extracted bone marrow was used directly to generate BMDC.

### Immature BMDC cultures

BMDC cultures were performed according to Alraies et al., 2024. Briefly, 1/3 of fresh total BM cell suspension was cultured in 100 mm TC-treated culture dishes (Corning, cat. #354450) at 37 °C, 5% CO2, and cells were replated at day 4 and 7. Passaging was performed using PBS-EDTA (5 mM) to detach cells and then replated in fresh media at a concentration of 0.5*10^6 cells/mL. Immature BMDCs (iDCs) were used at day 10, after recovering only the cells in suspension (adherent cells (granulocytes and macrophages) were discarded), when >90% of the cells were CD11c+ as per FACS validation.

### Mature BMDC cultures

Mature BMDC (mDC) cultures were obtained as described in Alraies et al., 2024. Briefly, fully developed iDCs were stimulated with 100 ng/mL Lipopolyssacaride (LPS) from Salmonella enterica serotype Typhimurium (Sigma, cat. #L6143) for 25 min at 37°C. Subsequently, LPS-containing media were removed by centrifugation (1300 rpm for 5 min), and cells were resuspended in fresh DC medium for three times, and re-plated overnight in the incubator. mDCs were used for experiments the following day.

### BMDC transfection

Second generation lentiviral particles were prepared by co-transfecting HEK-293T cells with the plasmid of interest (pMD2.G (Addgene, cat. #12259), psPAX2 (Addgene, cat. #12260) and pCDH-NLS-GFP (Addgene, cat. #132772), and packaging and envelope plasmids. 48 h after transfection, virus-containing supernatant was collected, centrifuged 300g for 5min to remove cell debris, and filtered through a 0.45 μm filter. Cleared supernatants were mixed with Lenti-X concentrator (Takara) and concentrated 100x, following the manufacturer’s instructions. 20 μL of concentrated lentiviral particles were used to infect 2 x 10^6^ BMDCs at day 5 in a 6-well plate. At day 8, cells were collected and washed 2x in sterile PBS to remove remaining viral particles and used for experiments.

### Fluorescence in situ hybridization (FISH) of BMDCs

Cells attached to coverslips were fixed with methanol:acetic acid (3:1) for 15 minutes at room temperature or overnight at −20°C, rinsed in 80% ethanol and air dried for 5 minutes. XMP Y Green (MetaSystems, cat. # D-1421-050-FI) and XMP X Orange (MetaSystems, cat. # D-1420-050-OR) probe mixtures were applied and sealed with a coverslip. Slides were denatured at 75°C for 2 minutes and incubated at 37°C overnight in a humidified chamber. Slides were washed with 0.4X SCC at 72°C for 2 minutes, 2X SCC, 0.1% Tween-20 at room temperature for 30 sec, and rinsed with PBS. Slides were incubated with DAPI solution for 10 minutes before mounting in anti-fade reagent. Images of ∼10 µm stack (with step size z = 0.5 µm) were acquired on an inverted laser scanning confocal Leica SP8 microscope at 63X oil immersion objective with numerical aperture 1.4. Images were analysed with Imaris software (version 9.8.2, Oxford Instruments) to quantify cells aneuploid for Y or X and total number of cells. Aneuploid cells were declared as cells with 0 or >1 copies of chromosome Y or X. Cells on the edges were excluded from the analysis.

### Microchannel device preparation and immunofluorescence analysis

Polydimethylsiloxane (PDMS) (Neyco, cat. #RTV 615) micro-channel chips were produced as previously described (Faure-André et al., 2008; Vargas et al., 2016). In this study, 5 μm x 5 μm channels stuck onto glass coverslips (FisherScientific, cat. #12323138) were used. The surface was coated with 10 μg/ mL bovine plasma fibronectin (Sigma, cat. #F1141) for 30 min at 37°C and washed with DC media. 0.1*10^6^ cells were loaded into access port in DC media. Cells were allowed to freely migrate inside the microchannels for 16h. Afterwards, the channels were washed with PBS and incubated with 4% paraformaldehyde (PFA, Pierce, cat. #28906) diluted in PBS for 30 min at RT. After fixation, samples were washed with PBS and the channels were gently separated from the cover. The glass cover with the attached cells was washed with PBS and immunofluorescence was proceeded as described above.

### Microchannel device preparation and NE-rupture analysis

Microchannels of different size (MC004, 4D Cell) were prepared according to the manufacturer’s protocol and coated with 10 μg/mL of fibronectin the day before the experiment. Cells were introduced in microchannels by plating 100.000 BMDCs on each access port at least 6 h before the experiments. Overnight live imaging (14 h) of NLS-GFP positive cells migrating inside microchannels were imaged on a Nikon Eclipse Ti-E inverted fluorescent microscope equipped with DC-152Q-C00-FI using NIS V4.30 software (Nikon). At least two fields per condition were acquired per experiment. Image analysis was carried out with Fiji ImageJ v.1.54f.

### Quantification of nuclear ruptures in migrating cells

Raw acquisitions were used to reconstruct individual videos, which were aligned and stabilized using MultiStackReg registration plugin in ImageJ. The percentage of nuclear rupture events was manually quantified by inspecting reconstructed videos of NLS-GFP positive cells migrating inside the channels. Migrating cell kymographs were generated by subtracting from each frame the mean projection of the whole movie, generating clear objects in a dark background.

### Calculation of immunofluorescence intensity

Fluorescence images were acquired as z stacks of 0.33 μm width. Intensity was calculated by a customized macro created in ImageJ (v2.9.0/1.53t, National Institutes of Health) that measured the mask signal of the nucleus (DAPI stained) and the whole cell (Phalloidin stained). Subsequently, the “analyze particles” tool was used to determine the area and elongation, and mean intensity of each mask.

### BMDC Electron Microscopy

BMDCs were prepared as described above and cells were let migrating overnight inside channels printed in a PDMS chip. For electron microscopy analysis, after PDMS removal, cells were fixed using 2.5% glutaraldehyde (Sigma, cat. # G5882) in 0.1 M cacodylate buffer (Electron Microscopy Sciences, cat. #11652), pH 7.4 for 1h, post fixed for 1h with 2% buffered osmium tetroxide (Electron Microscopy Sciences, cat. #19150), dehydrated in a graded series of ethanol solution, and then embedded in epoxy resin (EMbed 812 Embedding Kit, cat. #14121). Images were acquired with a digital 4k CCD camera Quemesa (EMSIS GmbH, Münster, Germany) mounted on a Tecnai Spirit transmission electron microscope (ThermoFisher, Eindhoven, Netherlands) operated at 80 kV.

### Pharmacological treatments on BMDC

2*10^6^ cells were incubated with 25μM CK666 (Tocris, cat. #3950) or 1μM Etoposide (Sigma-Aldrich, cat. #E1383) diluted in DC media. The samples were incubated from day 7 until day 10 (3 days of culture) or two hours at day 10 of culture at 37°C and 5% CO2. After treatment, cells were subjected to immunofluorescence staining procedures as described above.

### Image Streamer flow cytometry analysis - Staining protocol

Samples, single-color controls and unstained samples containing 2.5*10^6^ cells were washed with fresh ice-cold MACs Buffer (PBS-0.5%BSA-2mM EDTA). Subsequently, cells were subjected to Live/Dead staining (LIVE/DEAD™ Fixable Far Red - Dead Cell Stain Kit, Thermo Fischer, cat#L10120) for 15 min at RT in darkness. Then, cells were washed with MACs Buffer and centrifuged for 1 min at 1800 rpm. The pellet was blocked with Purified Rat Anti-Mouse CD16/CD32 and PE-eFluo610-CD11c (Table S4) and incubated for 10 min at 4°C in darkness. Next, cells were washed and centrifuged as described above and fixed using freshly prepared PFA 4% + 0.001%Triton100X for 10 min at RT. After washing, cells were stained with Hoechst 33258 Staining Dye Solution (Abcam, cat. #ab228550) for 45 min at 4°C in darkness. Finally, samples were washed and resuspended in MACs Buffer for acquisition.

### Image Streamer flow cytometry analysis - Sample acquisition

Samples were analyzed at the single-cell level using imaging flow cytometry (ImageStream X MKII, Amnis/Cytek). The acquisition template was set to detect CD11c in channel Ch04 with a 561 nm laser (100 mW), Hoechst in channel Ch07 with a 405 nm laser (80 mW) and Live/Dead in channel Ch11 with a 642 nm laser (130 mW). Ch01 and Ch09 were used for Brightfield imaging. Single-color controls and unstained samples were used to calculate the compensation matrix.

Analysis was performed using IDEAS software (v6.2). The first selection, “Focus,” was done using the Gradient RMS feature on Ch01 to select events in focus. Next, “Singlets” were selected based on the area and aspect ratio in Ch01. Dead cells were excluded based on the intensity in Ch11, and positive events for Hoechst were isolated using the Ch07 intensity (“Live Ho^+^”). The “Circu” step was used to exclude events with unusual shapes in Ch01 (feature Circularity, which measures the degree of deviation of an object from a perfect circle). CD11c-positive events were selected based on Ch04 intensity (“CD11c^+^”), and cell cycle analysis was performed using Ch07 intensity to differentiate the “G0/G1,” “S,” and “G2/M” phases.

In each phase, circularity and elongation of the Ch07 staining were analyzed (using the Elongatedness feature, which measures the ratio of height to width), employing a stringent mask on the nucleus (Morphology Mask M07, Ch07). Three gates were defined: “High Circu Low Elong,” “Low Circu High Elong,” and “Low Elong Low Circu.” For each gate, the Lobe Count feature (which measures the number of symmetry axes to determine the number of lobes) was applied with the Morphology Mask M07, Ch07 to isolate events with “1 Lobe,” “2 Lobes,” or “Multi Lobes.”

### Quantification of Arpc5 immunostaining in BMDCs

ArpC5 subcellular localization in wildtype BMDCs was quantified based on SIM micrographs upon immunostaining for ArpC5 and nuclear counterstain (DAPI). Per ROI, the percentage of cells that displayed Arpc5-positive nuclear foci was assessed.

### RNA-sequencing – Sample preparation

Total RNA extraction from BMDCs generated as described above was performed using the RNeasy Mini RNA kit (Qiagen) including on-column DNase digestion according to the manufacturer’s protocol. The integrity of the RNA was confirmed in BioAnalyzer using RNA 6000 Pico kit (Agilent Technologies) (8.8 < RNA integrity no. (RIN) < 10). RNA-seq libraries were prepared from 300 ng to 1 µg of total RNA using an Illumina TruSeq Stranded mRNA Library Preparation kit and an Illumina Stranded mRNA Prep Ligation kit (Illumina, cat# 20040532), which allowed strand-specific RNA-seq. First, poly(A) selection using magnetic beads was performed to focus sequencing on polyadenylated transcripts. After fragmentation, cDNA synthesis was performed, and the resulting fragments were used for dA-tailing and were ligated to the TruSeq indexed adapters (for the TruSeq kit) or RNA Index Anchors (for the mRNA Ligation kit). PCR amplification was performed to generate the final indexed cDNA libraries (with 13 cycles). Individual library quantification and quality assessment was performed using a Qubit fluorometric assay (Invitrogen) with a dsDNA High-Sensitivity Assay kit and LabChip GX Touch using a High-Sensitivity DNA chip (PerkinElmer). Libraries were then pooled at equimolar concentrations and quantified by qPCR using a KAPA library quantification kit (Roche). Sequencing was performed on a NovaSeq 6000 instrument (Illumina) using paired-end 2 × 100 bp sequencing with an average depth of 30 × 106 clusters per sample. RNAseq data have been deposited at GEO (accession GSE288480).

### RNA-sequencing – Data analysis

Genome assembly was based on the Genome Reference Consortium (mm10). The quality of the RNA-seq data was assessed using FastQC (v.0.11.9). Reads were aligned to the transcriptome using STAR (v.2.6.1). Genes with a low number of counts (<10) were filtered out. Differentially expressed genes (DEGs) analysis was performed using DESeq2 (v.1.22.2) with the design ‘∼ genotype’ for each time point (Day 7, 8, 9 and 10). DEGs were identified based on Padj < 0.05 and absolute log2(FC) > 0.5. GO Biological Process enriched in KO samples compared to WT for each day was done using EnrichR package (Kuleshov et al., 2001). For gene clustering, all DEGs with log2 fold change > 1 were concatenated and their expression was compared in samples from all time points. K-means clustering was performed using the pheatmap package (v.1.0.12). GSEA61 was performed in the normalized count matrices from DESeq2 using the GSEA software (v.4.0.3) with the default parameters, except for the number of permutations that were fixed at n = 1,000. Volcano plots were generated with EnhancedVolcano. Senescence signatures were taken from the AgingAtlas (2021), apoptosis signatures were taken from the Hallmark MSigDB database (The Molecular Signatures Database; https://www.gsea-msigdb.org/gsea/msigdb).

### Statistical analysis

Number of mice and experiments, and statistical tests are reported in each figure legend. Analyses were performed using GraphPad Prism 8 software. Statistical significance was calculated using Student’s t test (unpaired) or Mann-Whitney, One-way ANOVA, two-way-ANOVA and logistic regression (GLM: Generalized Linear Model tool, Prism software) with binomial distribution in the case of the aneuploidy data, according to test requirements. Error bars represent SEM or SD and p values <.05 were considered statistically significant (* p <.05, ** p <.01, *** p <.001, **** p <.0001).

## ACKNOWLEDGEMENTS

We thank Institut Curie for access to the flow cytometry and the cell and tissue imaging facilities (PICT-IBiSA). Moreover, we are grateful to the animal facilities at Institut Curie and Saarland University for expert service. We thank Evelyn Wirth for support in mouse genotyping. We thank Tim Lämmermann, Mathieu Piel, Sarah Lambert, Raphael Ceccaldi, Arnaud Echard and Iden lab members for their valuable comments on this work.

## DATA AVAILABILITY

Correspondence and requests for materials related to this study should be sent to. All data supporting the findings of this study are available within the manuscript and its supplemental information, or from the corresponding authors on reasonable request.

## AUTHOR CONTRIBUTIONS

Conceptualization: M.G.D., A.K.B., S.I., A.M.L.D.; Methodology: M.G.D., A.K.B., V.S.R., Z.A., D.F., F.B., C.G., F.G., N.M.; Validation: M.G.D., A.K.B., S.I., A.M.L.D.; Formal analysis: M.G.D., A.K.B., S.I., A.M.L.D.; Investigation: M.G.D., A.K.B., C.C.J., V.S.R., S.I., A.M.L.D., J.V., R.A., D.S., N.D.S., L.L.M., A.S., A.C., M.S.R., M.M., V.C., A.D., G.B.; Resources: A.M.L.D., S.I.; Writing: A.M.L.D., S.I.; Editing: M.G.D., A.K.B., S.I., A.M.L.D.; Visualization: M.G.D., A.K.B.; Supervision: S.I., A.M.L.D.; Project administration: A.M.L.D., S.I.; Funding acquisition: A.M.L.D., S.I.

