## Supplementary material for "The Arp2/3 complex maintains genome integrity and survival of epidermal Langerhans cells": suppl_info

**Supplementary Information Delgado, Burkhart et al.**

Inventory


- Supplementary Figure Legends
- Supplementary Figure 1
- Supplementary Figure 2
- Supplementary Figure 3
- Supplementary Figure 4
- Supplementary Figure 5
- Supplementary Figure 6
- Supplementary Figure 7
- Supplementary Table 1
- Supplementary Table 2
- Supplementary Table 3
- Supplementary Table 4
- Supplementary Table 5
- Supplementary Table 6
- Supplementary Table 7

### Supp. Figure 1, related to main Figure 1

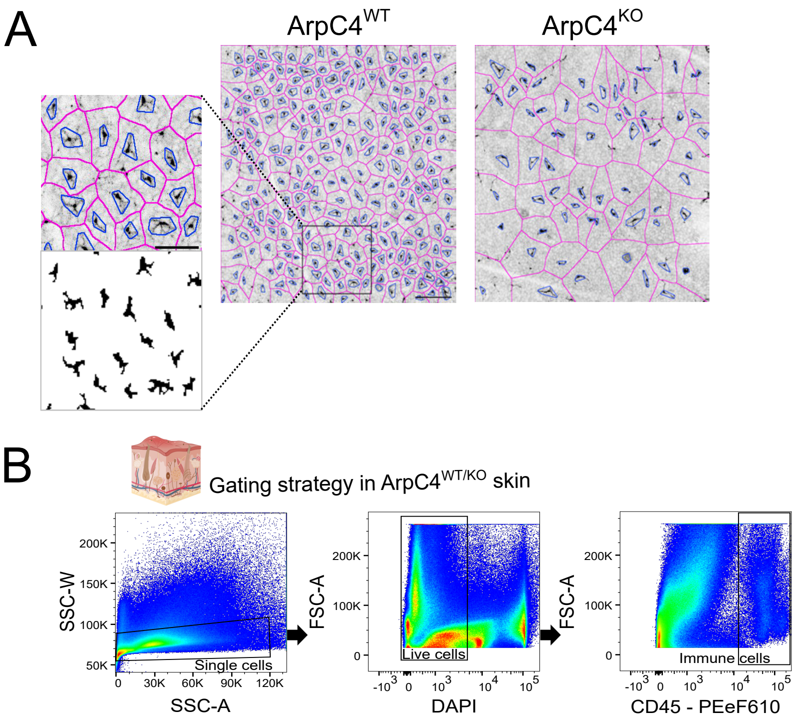

### Supp. Figure 2, related to main Figure 2

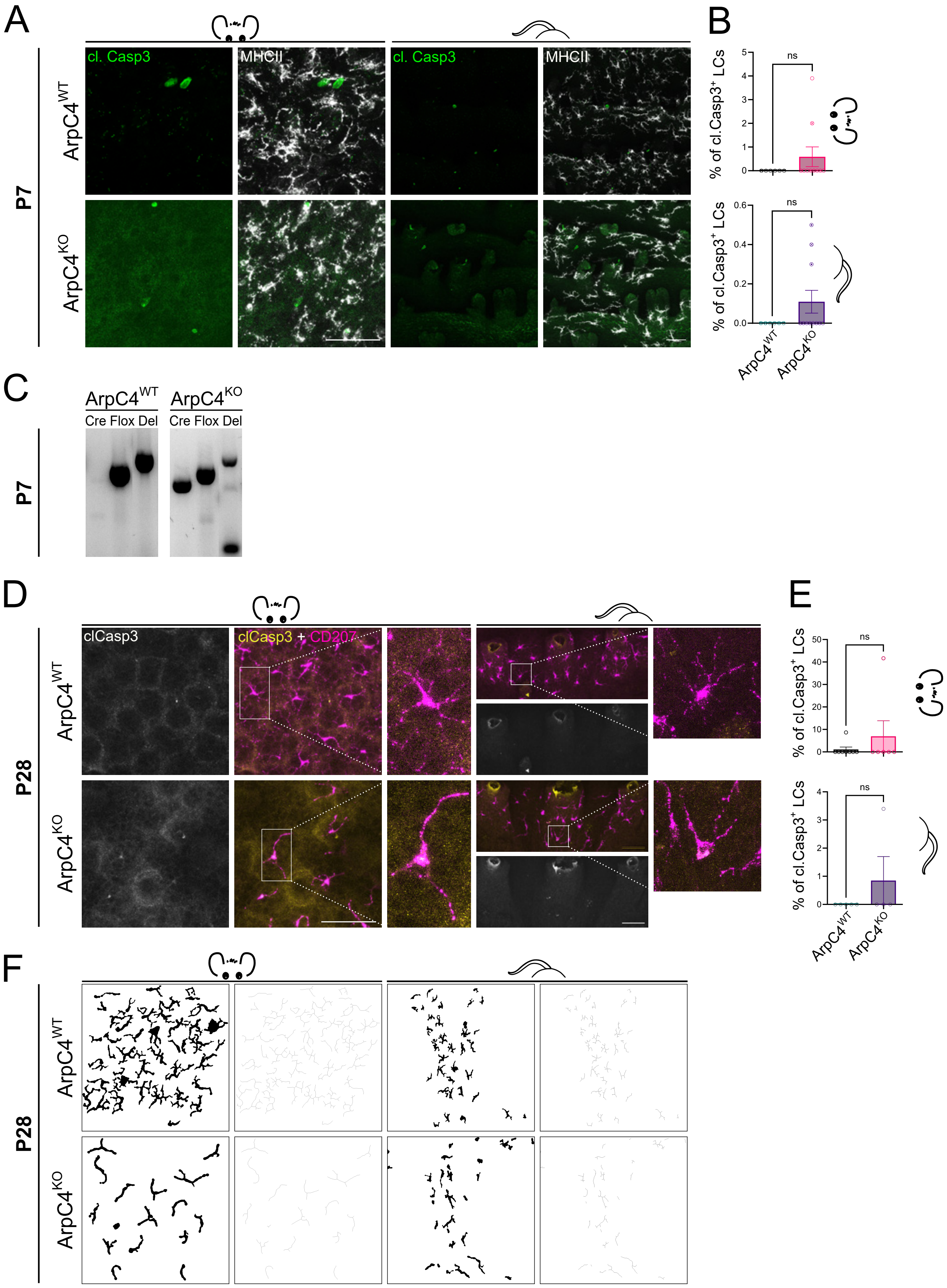

### Supp. Figure 3, related to main Figure 3

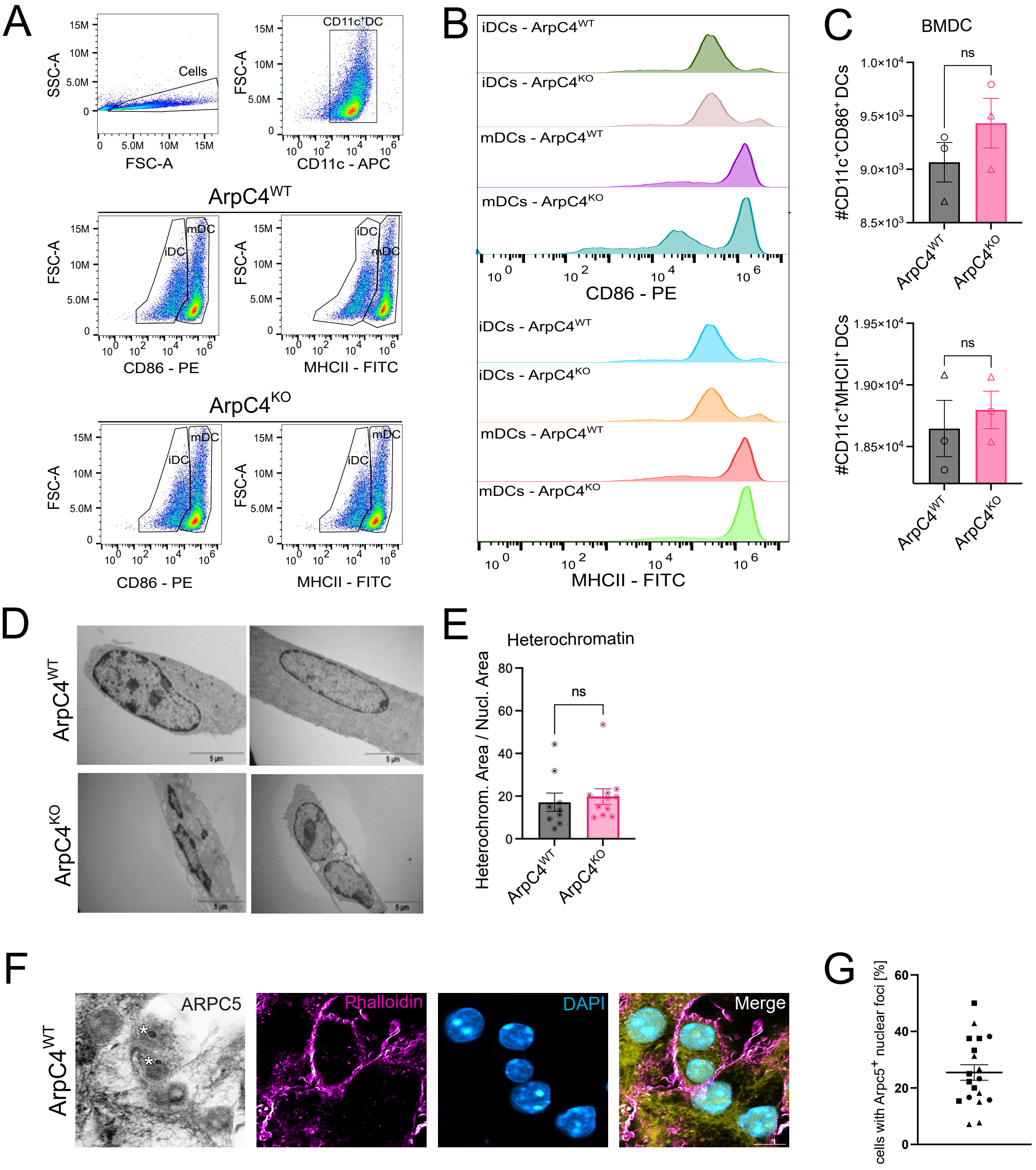

### Supp. Figure 4, related to main Figure 4

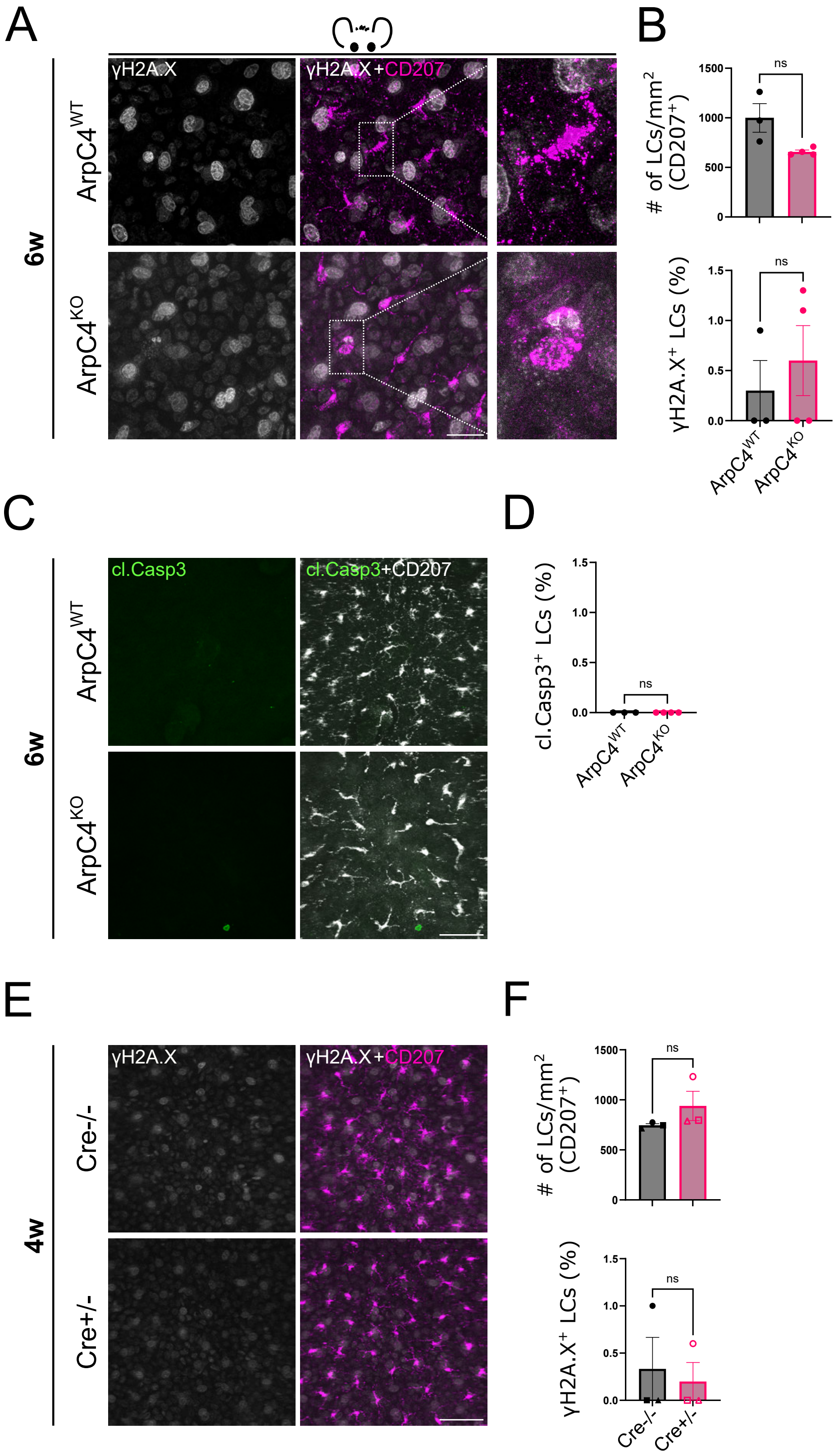

### Supp. Figure 5, related to main Figure 5

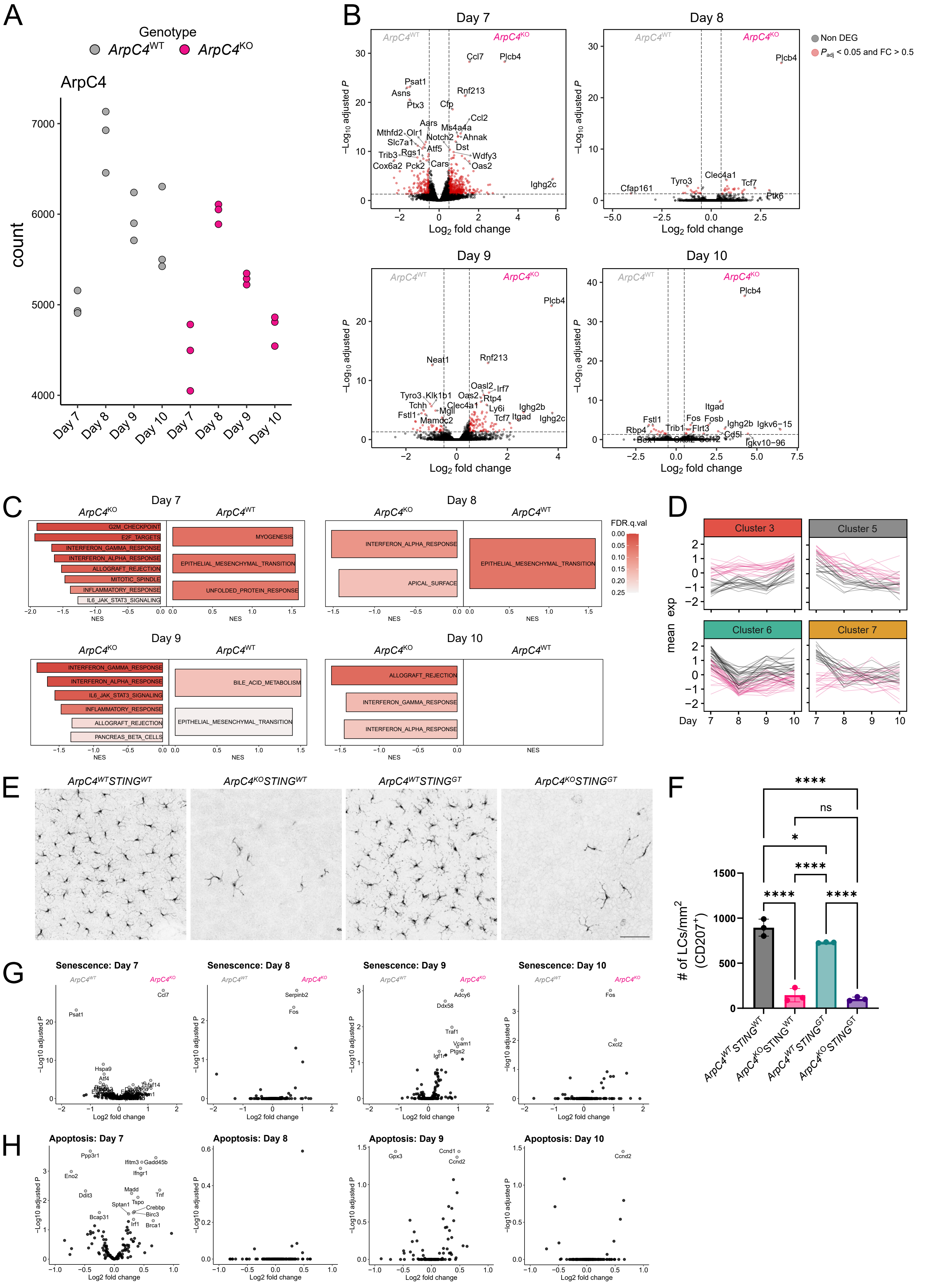

### Supp. Figure 6, related to main Figure 6

A

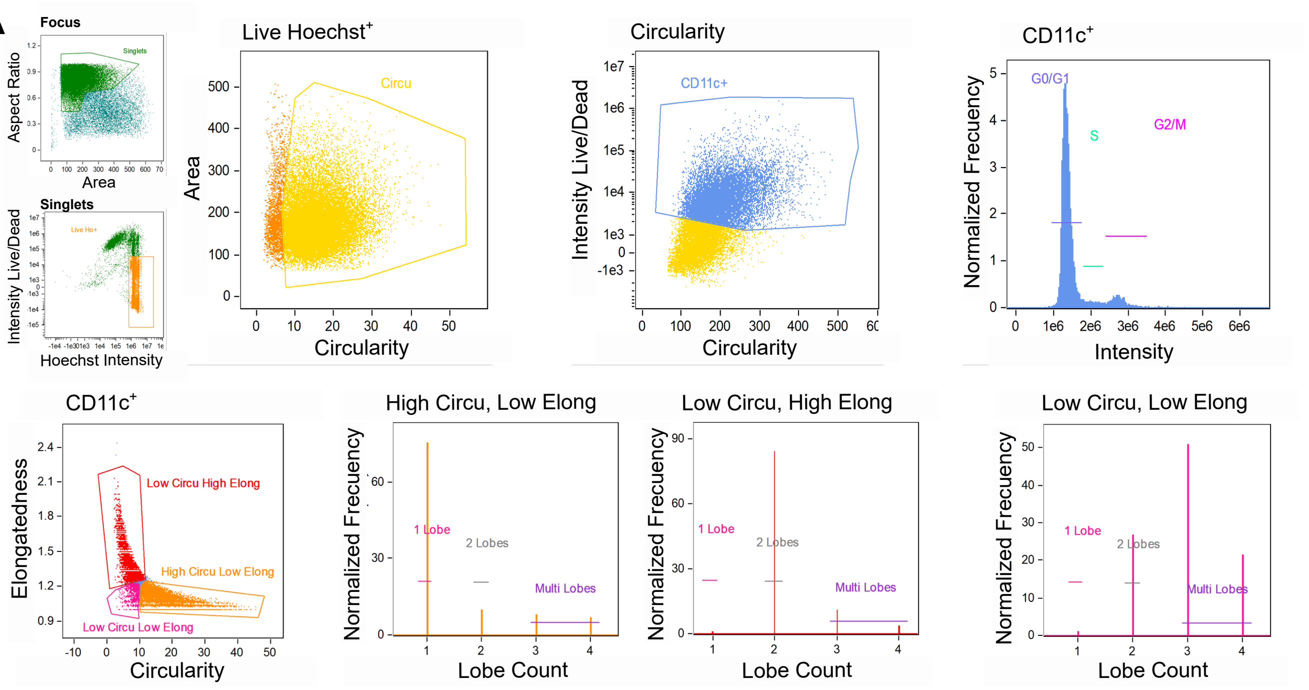

### Supp. Figure 7, related to main Figure 7

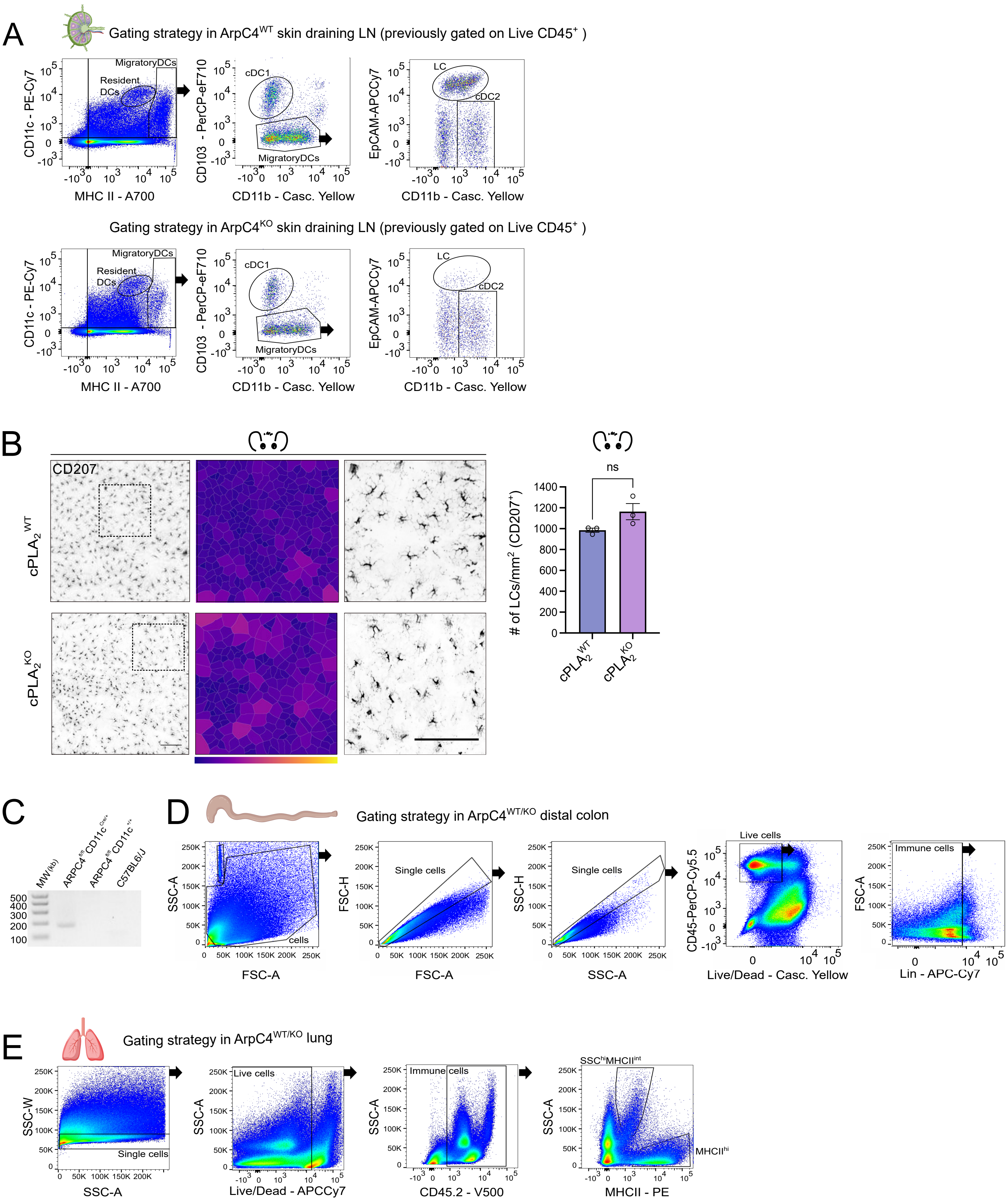

#### SUPPLEMENTARY FIGURE LEGENDS

**Supplementary Figure 1, related to main Figure 1:**
**Image segmentation for LC distribution in epidermis, and gating strategies**

(A) Representative image of the analysis performed in Figure 1 to generate Voronoi diagrams. $CD207^{+}$ signal was Gaussian blurred, segmented and used to generate a mask after thresholding the image. The signal was skeletonized using analysis tool Skeleton 2D/3D from ImageJ software. For each cell a convex value (blue line) and then a Voronoi (magenta line) was generated using the software tool.

(B) Gating strategy in skin; single cells (left panel) were used to identify live cells (DAPI-negative, center panel), then CD45 was used to identify immune cells (right panel).

**Supplementary Figure 2, related to main Figure 2:**
**Deletion of Arpc4 does not interfere with the initial establishment of postnatal LC networks**

(A) Micrographs of immunostainings for LCs ($MHCII^{+}$; grey) and cl. Casp3 (green) in P7 ear (left) and tail (right) epidermal sheets. Scale bars = 50 µm.

(B) Quantification of (A): percentage of cleaved Caspase 3-positive LCs in P7 ear (top) and tail (bottom) epidermis. Ear: (n = 6 ctrl, 10 KO mice; ns p = 0.5; mean ± SD; two-sided Mann-Whitney U-test). Tail: (n = 6 ctrl, 11 KO mice; ns p = 0.3529; mean ± SD; two-sided Mann-Whitney U-test).

(C) Agarose gel of PCRs to detect Cre and floxed Arpc4 alleles in genomic DNA samples of tail skin epidermis at P7 (Cre, Flox). The Del PCR specifically confirms the deletion of the floxed region in the Arpc4 gene, $ArpC4^{KO}$: $CD11cCre^{wt/+}$;$ArpC4^{fl/fl}$.

(D) Micrographs of immunostainings for LCs ($CD207^{+}$; magenta) and cleaved Caspase 3 (cl. Casp3; yellow) in ear (left) and tail (right) epidermal sheets of 4-weeks-old mice. Scale bars = 50 µm.

(E) Quantification of (D): percentage of cl. Casp3-positive LCs in ear (top) and tail (bottom) epidermis. Ear: (n = 8 ctrl, 7 KO mice; ns p = 0.5; mean ± SD; two-sided Mann-Whitney U-test). Tail: (n = 5 ctrl, 4 KO mice; ns p = 0.4444; mean ± SD; two-sided Mann-Whitney U-test). Scale bars = 50µm; cl. Casp3 = cleaved Caspase 3.

(F) Thresholded and skeletonized images of micrographs of ear (left panel) and tail (right panel) epidermal sheets of 4-week-old mice immune stained for CD207/langerin.

**Supplementary Figure 3, related to main Figure 3:**
**BMDC differentiation and heterochromatin analyses in control and $Arpc4^{KO}$ BMDCs.**

(A) LPS treatment was used to activate immature DCs (iDCs), resulting in the generation of a mature population (mDCs). Gating strategy used; single $CD11c^{+}$ cells were stained with activation markers; CD86 and MHCII expression serves to discriminate the immature (iDC) and mature (mDC) population in ctrl (upper panel) and $ArpC4^{KO}$ cells (lower panel).

(B) Histograms for each population show a shift of the level of expression of CD86 (upper panels) and MHCII (lower panels).

(C) Number of $CD11c^{+}$;$CD86^{+}$ DCs in ctrl and $ArpC4^{KO}$ BMDCs (top panel; ns p = 0.2844; mean ± SD; unpaired, two-sided t-test) and number of $CD11c^{+}$;$MHCII^{+}$ DCs in MHCII expression (bottom panel; ns p = 0.6105; mean ± SD; unpaired, two-sided t-test).

(D) Representative electron microscopy images of ctrl (upper panels) and $ArpC4^{KO}$ (lower panel) BMDCs inside 8 µm × 5 µm fibronectin-coated µchannels. Scale bars = 5µm.

(E) Quantification of heterochromatin inside whole nuclear area of ctrl and $ArpC4^{KO}$ cells, n ctrl = 9, KO = 11 cells, ns p = 0.6402; mean ± SD; unpaired, two-sided t-test.

(F) Exemplary super-resolution SIM images of wild-type BMDCs immunostained for ArpC5 (grey), phalloidin (magenta), DAPI (cyan), and merge. Asterisks mark Arpc5-positive foci in the nucleus. Scale bar = 5µm.

(G) Quantification of (F) regarding Arpc5 immunostaining in nuclear foci of control cells. n = 3 mice (each symbol represents a biological replicate, while each dot represents the value per ROI).

**Supplementary Figure 4, related to main Figure 4:**

**DNA damage and apoptosis in control and *ArpC4*$^{KO}$ LCs *in vivo*.**

(A) Micrographs of immunostainings for LCs (CD207$^{+}$; magenta) and γH2Ax (white) in ear epidermal sheets of 6-weeks-old mice. Scale bar = 20 µm.

(B) Quantification of (A): (top) LC density (LC number/mm²) in ear epidermis (n = 3 ctrl, 4 KO mice; ns p = 0.1401; mean ± SD; unpaired two-tailed Student's t-test); (bottom) percentage of γH2Ax-positive LCs in ear epidermis (ns p = 0.5429; mean ± SD; two-sided Mann-Whitney U-test).

(C) Micrographs of immunostainings for LCs (CD207$^{+}$; white) and cl. Casp3 (green) in ear epidermal sheets of 6-weeks-old mice. Scale bar = 50 µm.

(D) Quantification of (C): percentage of cl. Casp3-positive LCs in ear epidermis (n = 3 ctrl, 4 KO mice; ns p>0.9999; mean ± SD; two-sided Mann-Whitney U-test).

(E) Micrographs of immunostainings for LCs (CD207$^{+}$; magenta) and γH1A.X (white) in ear epidermal sheets of 4-weeks-old CD11c-Cre$^{+/-}$ and CD11c-Cre$^{-/-}$ mice. Scale bar = 50 µm.

(F) Quantification of (E): (top) LC density (LC number/mm²) in ear epidermis (n = 3 mice; ns p = 0.1000; mean ± SD; two-sided Mann-Whitney U-test); (bottom) percentage of γH2Ax-positive LCs in ear epidermis (ns p>0.9999; mean ± SD; two-sided Mann-Whitney U-test).
cl. Casp3 = cleaved Caspase 3

**Supplementary Figure 5, related to main Figure 5:**
**RNA sequencing of *in vitro* cultured control and *ArpC4*$^{KO}$ DCs.**

(A) RNAseq of control (black) and KO (magenta) BMDCs after 7, 8, 9, or 10 days of culture; ArpC4 normalized expression per condition.

(B) Volcano plots of genes differentially expressed between ArpC4$^{WT}$ and ArpC4$^{KO}$ at the indicated days. Red points are genes with adjusted p value < 0.05 and log2 fold change > 0.5.

(C) GSEA analysis between ArpC4$^{WT}$ and ArpC4$^{KO}$ DCs at each indicated day in culture. Enriched pathways with FDR<0.25 are shown.

(D) Expression of genes over time. Genes are grouped by cluster (from Figure 5D).

(E) Micrographs of ear epidermal sheets from adult (11 weeks old) control (ArpC4$^{WT}$STING$^{WT}$), ArpC4$^{KO}$ (ArpC4$^{KO}$STING$^{WT}$), STING$^{KO}$ (ArpC4$^{WT}$STING$^{GT}$) and double KO (ArpC4$^{KO}$STING$^{GT}$) mice immunostained for LCs (CD207$^{+}$). Scale bar = 50 µm.

(F) Quantification of (E): LC density (LC number/mm²) in ear epidermis (n = 3 mice; **** < 0.0001; control vs. STING$^{KO}$: *p = 0.044; ArpC4$^{KO}$ vs. double KO: ns p = 0.837; Two-way ANOVA/Tukey's multiple t-test; mean ± SD).

(G) Volcano plots displaying differential expression between ArpC4$^{WT}$ and ArpC4$^{KO}$ of genes belonging to the Aging Atlas senescence signature at each time point. Significantly regulated genes are shown in gray and labeled.

(H) Volcano plots displaying differential expression between ArpC4$^{WT}$ and ArpC4$^{KO}$ of genes belonging to the Hallmark MSigDB mouse apoptosis signature at each time point. Significantly regulated genes are shown in gray and labeled.

**Supplementary Figure 6, related to main Figure 6:**
**Gating strategy for analysis of control and *ArpC4*$^{KO}$ BMDC on FACStreamer.**

(A) Analysis performed on the software IDEAS that integrates images with flow cytometry information. Only cells in focus were considered for the analysis and were gated on high aspect ratio (circularity and elongation) and low area. Afterwards, a single live (low intensity of Live/dead marker) and high Hoechst intensity population was identified. Then, cells were gated according to their area and circularity, and subsequently only CD11c$^{+}$ cells were selected for further analysis. In this population, cell cycle phases G0/G1, S and G2/M were analyzed as shown in the upper right panel. In this same population, elongation vs. circularity was visualized to distinguish three populations that were used to define the lobe count: (1) High circularity, low elongation describing the single lobe population, (2) Low circularity, high elongation describing the double-lobe population and (3) Low circularity, low elongation that describes the multi-lobe population. The normalized frequency of these populations is shown on the graphs to the right.

**Supplementary Figure 7, related to main Figure 7:**
**Gating strategies for analysis of control and *ArpC4*$^{KO}$ myeloid and monocyte-derived cell populations *in vivo.***

(A) Gating strategy for lymph node samples: single live cells were used to identify resident and migratory DCs according to their expression of CD11c and MHCII. Then, in the migratory DCs, two populations were defined according to their level of expression of CD103 (low: migratory DCs, and high: cDC1s). Within the gate of migratory DCs, LCs were identified as EpCAM$^{high}$, whereas cDC2s were CD11b$^{high}$, EpCAM$^{low}$.

(B) Micrographs of ear epidermal sheets from adult (14 weeks old) control (cPLA$_2^{WT}$) and cPLA$_2^{KO}$ mice immunostained for LC marker (CD207/langerin). Middle: Voronoi diagram of LCs generated from micrographs. Scale bar = 100 µm. Right micrograph: magnification of marked areas. Scale bar = 100 µm. Right: Quantification of LC density (LC number/mm²) in ear epidermis (N= 1 experiment/ 6 mice; 3 Ctrl and 3 KO; ns p = 0.0896; mean ± SD; unpaired two-tailed Student's t-test).

(C) Genotyping PCR of colon macrophages after CD11c-Cre-mediated deletion in the ARPC4 gene (174 bp) plus Cre-negative and C57BL6/J controls

(D) Gating strategy for distal colon; single live CD45$^{+}$ cells (left panels) were used to identify lin-negative immune cells (right panels).

(E) Gating strategy for lung; single cells (left panels) were used to identify live CD45.2$^{+}$ immune cells, then two populations were identified according to their high granularity and intermediate MHCII expression level (SSC$^{hi}$MHCII$^{int}$) and high level of MHCII expression (right panel).

#### SUPPLEMENTARY TABLES

**Supplementary Table 1.** ***In vivo*** **analysis of DC subsets in skin and skin draining LNs**

**LNs**

| **Antibodies** | **Clone** | **SOURCE** | **IDENTIFIER** | **Dilution** |
|---|---|---|---|---|
| PE-Cyanine7, anti-CD11c | N418 | eBioscience | Cat# 25-0114-81 | 1:800 |
| APC/Cyanine7, anti-mouse CD326 (Ep-CAM) Antibody | G8.8 | Biolegend | Cat# 118217 | 1:800 |
| PE, Rat Anti-Mouse CD86 | GL1 (RUO) | BD Pharmingen | Cat# 553692 | 1:800 |
| Brilliant Violet 605 anti-mouse/human CD11b Antibody | M1/70 | Biolegend | Cat# 101237 | 1:500 |
| Alexa Fluor 700, MHC Class II (I-A/I-E) Monoclonal Antibody | M5/114.15.2 | eBioscience | Cat# 56-5321-80 | 1:250 |
| APC Rat Anti-Mouse CD8a | 53-6.7 | BD Pharmingen | Cat# 553035 | 1:100 |
| PerCP-eFluor 710 anti-CD103 (Integrin alpha E) Monoclonal Antibody | 2E7 | eBioscience | Cat# 46-1031-80 | 1:100 |

| **Isotype Control** | **Clone** | **SOURCE** | **IDENTIFIER** | **Dilution** |
|---|---|---|---|---|
| PE-Cyanine7, Armenian Hamster IgG | eBio299Arm | eBioscience | Cat# 25-4888-82 | 1:750 |
| APC/Cyanine7 Rat IgG2a, k | TK2758 | Biolegend | Cat# 400524 | 1:750 |
| PE, Rat LOU, also known as Louvain, LOU/C, LOU/M IgG2a, k | R35-95 | BD Pharmingen | Cat# 557229 | |
| Brilliant Violet 605™ Rat IgG2b, k | RTK4530 | Biolegend | Cat# 400650 | 1:500 |
| Alexa Fluor 700, Rat IgG2b kappa | eB149/10H5 | eBioscience | Cat# 56-4031-80 | 1:250 |
| APC, Rat LOU, also known as Louvain, LOU/C, LOU/M IgG2a, k | R35-95 | BD Pharmingen | Cat# 551139 | 1:100 |
| PerCP-eFluor 710, Armenian Hamster IgG | eBio299Arm | eBioscience | Cat# 46-1031-80 | 1:50 |

**Skin**

Additionally, for the skin, the following antibody/isotype control was added

| | | | | |
|---|---|---|---|---|
| PE-eFluor 610 anti-CD45 Monoclonal Antibody | 30-F11 | eBioscience | Cat# 61-0451-82 | 1:100 |
| PE-eFluor™ 610, Rat IgG2b k, Isotype Control | eB149/10H5 | eBioscience | Cat# 61-4031-80 | 1:250 |

**Supplementary Table 2. *In vivo* analysis of DC subsets in lung**

| Antibodies | Clone | SOURCE | IDENTIFIER | Dilution |
|---|---|---|---|---|
| V500, Mouse anti-Mouse CD45.2 | 104 | BD Horizon | Cat# 562129 | 1:100 |
| PE, Mouse Anti-Mouse I-A[b] | AF6-120.1 | BD Pharmingen | Cat# 553552 | 1:50 |
| PECy7, Anti-Mouse CD11c | N418 | Invitrogen | Cat# 25-0114-82 | 1:200 |
| PerCP-Cy5.5, Anti-Mouse CD11b | M1/70 | Invitrogen | Cat# 45-0112-82 | 1:200 |
| eFluor450, Anti-Mouse CD103 (Integrin alpha E) | 2E7 | Invitrogen | Cat# 48-1031-82 | 1:100 |
| Alexa700, Rat Anti-Mouse CD24 | M1/69 | BD Biosciences | Cat# 564237 | 1:50 |
| APC, anti-mouse CD64 (FcγRI) | X54-5/7.1 | Biolegend | Cat# 139306 | 1:100 |
| PE-CF594 Rat Anti-Mouse Siglec-F | E50-2440 | BD Horizon | Cat# 562757 | 1:200 |
| FITC, F4/80 Monoclonal Antibody | BM8 | Invitrogen | Cat# 11-4801-82 | 1:100 |

| Isotype Control | Clone | SOURCE | IDENTIFIER | Dilution |
|---|---|---|---|---|
| V500, Mouse IgG2a, k | G155-178 | BD Horizon | Cat# 561221 | 1:200 |
| PE, Mouse IgG2a, k | G155-178 | BD Pharmingen | Cat# 553457 | 1:200 |
| PE-Cy7, Armenian Hamster IgG | eBio299Arm | Invitrogen | Cat# 25-4888-82 | 1:200 |
| PerCP-Cy5.5, Rat IgG2b k | eB149/10H5 | Invitrogen | Cat# 45-4031-80 | 1:200 |
| eFluor450, Armenian Hamster IgG | eBio299Arm | Invitrogen | Cat# 48-4888-82 | 1:100 |
| Alexa700, Rat IgG2b, k | A95-1 | BD Pharmingen | Cat# 557964 | 1:100 |
| APC, Mouse IgG1, κ | MOPC-21 | Biolegend | Cat# 400120 | 1:200 |
| PE-CF594, Rat IgG2a, k | R35-95 | BD Horizon | Cat# 562302 | 1:200 |
| FITC, Rat IgG2a k | eBR2a | Invitrogen | Cat# 11-4321-80 | 1:200 |

**Supplementary Table 3.** ***In vivo*** **analysis of DC subsets in colon**

| Antibodies | Clone | SOURCE | IDENTIFIER | Dilution |
|---|---|---|---|---|
| PE-Cyanine5.5, CD45 Monoclonal Antibody | 30-F11 | eBioscience | Cat# 35-0451-82 | 1:500 |
| APC/Cyanine7, anti-mouse CD3ε Antibody | 145-2C11 | Biolegend | Cat# 100330 | 1:200 |
| PE-Cyanine7, CD11b Monoclonal Antibody | M1/70 | eBioscience | Cat# 25-0112-82 | 1:200 |
| PE, CD103 (Integrin alpha E) Monoclonal Antibody | 2E7 | eBioscience | Cat# 12-1031-82 | 1:200 |
| Brilliant Violet 421™ anti-mouse CD64 (FcγRI) Antibody | X54-5/7.1 | Biolegend | Cat# 139309 | 1:10 |
| PE/Cyanine5 anti-mouse F4/80 Antibody | BM8 | Biolegend | Cat# 123112 | 1:20 |
| PE/Dazzle™ 594 anti-mouse Ly-6C Antibody | HK1.4 | Biolegend | Cat# 128043 | 1:333 |
| Alexa Fluor 700, anti-mouse I-A/I-E Antibody | M5/114.15.2 | Biolegend | Cat# 107622 | 1:400 |
| APC Hamster Anti-Mouse CD11c | HL3 (RUO) | BD Pharmingen | Cat# 550261 | 1:10 |
| PerCP5.5 anti-CD45.5 | 104 (RUO) | BD | Cat# 552950 | 1:300 |
| APC anti-CD64 | X54-5/7.1 | Biolegend | Cat# 139306 | 1:200 |
| Brilliant Violet 650 anti-F4/80 | BM8 | Biolegend | Cat# 123149 | 1:200 |
| PE-Cy7 anti-CD11c | N418 | eBioscience | Cat# 25-0114-82 | 1:200 |

**Isotypes**

| | | | | |
|---|---|---|---|---|
| PE-Cyanine5.5, Rat IgG2b kappa | eB149/10H5 | eBioscience | Cat# 35-4031-80 | 1:200 |
| APC/Cyanine7 Armenian Hamster IgG | HTK888 | Biolegend | Cat# 400927 | 1:200 |
| PE-Cyanine7, Rat IgG2b kappa | eB149/10H5 | eBioscience | Cat# 25-4031-82 | 1:200 |
| PE, Armenian Hamster IgG | eBio299Arm | eBioscience | Cat# 12-4888-81 | 1:200 |
| Brilliant Violet 421™ Mouse IgG1, κ | MOPC-21 | Biolegend | Cat# 400158 | 1:200 |
| PE/Cyanine5 Rat IgG2a, κ | RTK2758 | Biolegend | Cat# 400509 | 1:200 |
| PE/Dazzle™ 594, Rat IgG2c, κ | RTK2758 | Biolegend | Cat# 400558 | 1:200 |
| Alexa 700, Rat IgG2b, κ | eB149/10H5 | eBioscience | Cat# 56-4031-80 | 1:200 |
| APC Hamster IgG1, λ1 | G235-2356 | BD Pharmingen | Cat# 553956 | 1:200 |

**Supplementary Table 4. FACs Streamer**

| Antibody | Clone | SOURCE | IDENTIFIER | Dilution |
|---|---|---|---|---|
| PE-eFluor 610, Anti-CD11c Monoclonal Antibody | N418 | eBioscience | Cat# 61-0114-82 | 2.5:200 |

**Supplementary Table 5A. Primary antibodies used for immunofluorescence of BMDCs**

| Antibody | SOURCE | IDENTIFIER | Dilution |
|---|---|---|---|
| Anti-Lap2B | BD Transduction Laboratories | Cat# 611000 | 1:1000 |
| Anti-Lamin B1 | Abcam | Cat# 16048 | 1:200 |
| Anti-Lamin A/C | Merck | Cat# SAB4200236 | 1:250 |
| Anti-phospho-Histone H2A.X (Ser139) | Merck | Cat# 05-636 | 1:200 |
| Anti-53pb1 | Invitrogen | Cat# PA1-16566 | 1:100 |
| Anti-RPA70/RPA1 Antibody | Cell Signaling Technologies | Cat# 2267 | 1:50 |
| Anti-ATR | Santa Cruz Biotechn. | Cat# sc-515173 | 1:50 |
| Anti-p16-ARC (ArpC5) | Synaptic Systems | Cat# 305011 | 1:100 |
| Anti-ds DNA antibody [35I9 DNA] | Abcam | Cat # ab27156 | 1:500 |

**Supplementary Table 5B. Secondary antibodies used for immunofluorescence of BMDCs**

| Antibody | SOURCE | IDENTIFIER | Dilution |
|---|---|---|---|
| Alexa 488, Goat, Anti-Rabbit Fab2 | Invitrogen | Cat# A11070 | 1:200 |
| Alexa 488, Goat, Anti-Mouse Fab2 | Invitrogen | Cat# A11017 | 1:200 |
| Alexa 546, Goat, Anti-Rabbit Fab2 | Invitrogen | Cat# A21202 | 1:200 |
| Alexa 546, Goat, Anti-Mouse Fab2 | Invitrogen | Cat# A11018 | 1:200 |
| Alexa 647, Goat, Anti-Rabbit Fab2 | Invitrogen | Cat# A21244 | 1:200 |
| Alexa 647, Goat, Anti-Mouse Fab2 | Invitrogen | Cat# A21236 | 1:200 |
| Phalloïdine Alexa Fluor™ 488 | Invitrogen | Cat# A12379 | 1:500 |
| Phalloïdine Alexa Fluor™ 546 | Invitrogen | Cat# A22283 | 1:500 |
| Phalloïdine Alexa Fluor™ 647 | Invitrogen | Cat# A22287 | 1:500 |

**Supplementary Table 5C. Secondary antibodies used for STED microscopy**

| Antibody | SOURCE | IDENTIFIER | Dilution |
|---|---|---|---|
| STAR RED, Goat anti-mouse IgG | Abberior | Cat# STRED | 1:200 |
| STAR ORANGE, Goat anti-rabbit IgG | Abberior | Cat# STORANGE | 1:200 |

**Supplementary Table 6A. Primary antibodies used for immunofluorescence stainings of tissues**

| Antibody | Clone | SOURCE | IDENTIFIER | Dilution |
|---|---|---|---|---|
| Rat IgG2b kappa monoclonal anti-I-A/I-E (MHCII) | M5/114.15.2 | BioLegend | Cat#107602; RRID: AB_313317 | 1:100 |
| Rat IgG monoclonal anti-CD207 (Langerin) | eBioRMUL.2 | eBioscience | Cat#14-2073-80; RRID: AB_493943 | 1:100 |
| Rabbit IgG polyclonal anti-active Caspase-3 | | R&D Systems | Cat#AF835 | 1:100 |
| Rabbit IgG monoclonal anti-Phospho-Histone H2A.X (Ser139) | clone 20E3 | Cell Signaling Technology | Cat#9718 | 1:200 |
| Rabbit IgG monoclonal anti-Lamin B1 | 702972 | Invitrogen™ | Cat#702972; RRID: AB_2784553 | 1:200 |
| Rabbit IgG polyclonal anti-Ki67 | | Abcam | Cat#ab15580 | 1:200 |

**Supplementary Table 6B. Primary antibodies used for immunofluorescence stainings of tissues**

| Antibody /Reagent | SOURCE | IDENTIFIER | Dilution |
|---|---|---|---|
| Alexa Fluor 488, Donkey, anti-Rat | Invitrogen | Cat# A21208 | 1:500 |
| Alexa Fluor 647, Goat, Anti-Rat | Invitrogen | Cat# A21247 | 1:500 |
| Alexa Fluor 647, Goat, Anti-Rabbit | Invitrogen | Cat# A11036 | 1:500 |
| Alexa Fluor 568, Goat, Anti-rabbit | Invitrogen | Cat# A21244 | 1:500 |
| DAPI | Carl Roth | Cat# 6335.1 | 1:500 |

**Supplementary Table 7.**

| Genotyping PCR | Oligo | Sequence |
|---|---|---|
| CD11c-Cre (Cre: 750 bp) | CD11c_Fw | GAC AAC TTC CCT CCT GGT CTC TG |
| | Cre_Rv | CCC AGA AAT GCC AGA TTA CG |
| ArpC4-Flox (wt: 620 bp; flox: 850 bp) | ArpC4_39529_F | AAG CCT TGC CCG AGA TAA TG |
| | ArpC4_39529_R | AAG CAA AGC CAG TCC CTC AC |
| ArpC4-deletion (del: 174 bp, flox 980bp) | tm1c_F | AAG GCG CAT AAC GAT ACC AC |
| | Floxed_LR | ACT GAT GGC GAG CTC AGA CC |

Cycling protocol

| Step | Temperature | Time | Cycles |
|---|---|---|---|
| Initial denaturation | 95°C | 5 min | 1 |
| Denaturation | 95°C | 45 sec | 34 |
| Hybridization | 51°C | 30 sec | |
| Extension | 72°C | 1 min | |
| Final extension | 72°C | 10 min | 1 |
